# Metabolic Alterations in Human Peripheral Blood Mononuclear Cells Associate with Progression to Islet Autoimmunity and Type 1 Diabetes

**DOI:** 10.1101/658500

**Authors:** Partho Sen, Alex M. Dickens, María Asunción López-Bascón, Tuomas Lindeman, Esko Kemppainen, Santosh Lamichhane, Tuukka Rönkkö, Jorma Ilonen, Jorma Toppari, Riitta Veijola, Heikki Hyöty, Tuulia Hyötyläinen, Mikael Knip, Matej Orešič

## Abstract

Previous metabolomics studies suggest that type 1 diabetes (T1D) is preceded by specific metabolic disturbances. Here we asked whether distinct metabolic patterns occur in peripheral blood mononuclear cells (PBMCs) of children later developing pancreatic *β*-cell autoimmunity or overt T1D. In a longitudinal cohort setting, PBMC metabolomic analysis was applied in children who either (1) progressed to T1D (PT1D, n=34), (2) seroconverted to ≥1 islet autoantibody without progressing to T1D (P1Ab, n=27), or (3) remained autoantibody negative during follow-up (CTRL, n=10). During the first year of life, levels of most lipids and polar metabolites were lower in PT1D and P1Ab, versus CTRLs. Pathway overrepresentation analysis suggested alanine, aspartate, glutamate, glycerophospholipid and sphingolipid metabolism were overrepresented in PT1D. Genome-scale metabolic models of PBMCs in T1D progression were developed using available transcriptomics data and constrained with metabolomics data from our study. Metabolic modeling confirmed altered ceramide pathways as specifically associated with T1D progression.

## INTRODUCTION

Type 1 diabetes (T1D) is a chronic, immune-mediated disease characterized by selective loss of insulin-producing β-cells in the pancreatic islets of genetically susceptible individuals (Atkinson and Eisenbarth, 2001; Knip et al., 2017). Over the past few decades, the incidence of T1D in most Western countries has been increasing, particularly among children below the age of five (Mayer-Davis et al., 2017). About 70% of children with T1D carry increased risk-associated genotypes in human leucocyte antigen (HLA) loci. On the other hand, only 3-7% of the population with the same risk alleles develop T1D (Achenbach et al., 2005).

The onset of clinical T1D is preceded by an asymptomatic prodromal period. The appearance of autoantibodies against insulin (IAA), an 65 kD isoform of glutamic acid decarboxylase (GADA), the insulinoma-associated antigen (IA-2A), and/or zinc transporter 8 (ZnT8A) in the plasma is an early sign for emerging islet autoimmunity and clinical T1D (Knip et al., 2017). It is known that especially children with multiple islet autoantibodies have an increased risk of T1D (Ziegler et al., 2013). In addition to genetic predisposition, other exogenous environmental factors affect risk, such as intestinal dysbiosis, reduced gut microbial diversity (Kostic et al., 2015), level of hygiene (Vatanen et al., 2016) and infant feeding regimen (Knip et al., 2010) are implicated in the initiation of β-cell autoimmunity. Our recent data also suggest that prenatal exposure to environmental chemicals modulates lipid metabolism in newborn infants and increases their subsequent risk of T1D (McGlinchey et al., 2019). However, the early pathogenesis of T1D is still poorly understood and the identification of molecular signatures and related pathways predictive of progression to overt T1D is still an unmet medical need.

Given our current level of understanding, potential alterations in immune cell metabolism may affect the host immune system (Buck et al., 2015). In fact, external perturbation of key metabolic processes such as glycolysis and amino acid metabolism have already been shown to impair T-cell activation, differentiation and cytokine production (Almeida et al., 2016; Berod et al., 2014; Geiger et al., 2016; Ma et al., 2017). Human peripheral blood mononuclear cells (PBMCs), including T cells (~70%), B cells (~15%), monocytes (~5 %), dendritic cells (~1%) and natural killer (NK) cells (~10%) obtained from healthy donors and T1D progressors are already being investigated in order to better understand this phenomenon (Sen et al. 2018). Such efforts seek to elucidate how immune cell metabolic processes are altered in seroconversion and progression to overt T1D; currently a largely unknown area.

Metabolomics is the study of small (< 1500 Dalton) molecules and their functions in cells, tissues and body fluids (Hollywood et al., 2006). The metabolome, which can be seen partly as a phenotypic readout of the genome, is sensitive to changes in immune system status, diet and gut microbiota (Frohnert and Rewers, 2016; Holmes et al., 2008; Oresic, 2012). Through metabolomic analyses, we have previously shown that decreased levels of plasma sphingomyelins (SMs) and phosphatidylcholines (PCs) are associated with progression to T1D (La Torre et al., 2013; Lamichhane et al., 2018a; Oresic et al., 2013; Oresic et al., 2008).

Here, we applied metabolomics to determine levels of molecular lipids and polar metabolites in PBMCs isolated from prospective samples collected in the Type 1 Diabetes Prediction and Prevention (DIPP) study, with the aim of elucidating the events preceding the onset of islet autoimmunity and overt T1D. We sought to address whether distinct metabolic patterns can be discerned during infancy among three study groups of children: (1) those, who developed clinical T1D, (2) those, who seroconverted to at least one islet autoantibody but were not diagnosed with T1D during follow-up, and (3) controls subjects, *i.e.*, children who remained autoantibody negative during follow-up.

## RESULTS

### Global lipidome of immune cells in progression to islet autoimmunity and T1D

PBMCs were isolated from children who (1) progressed to clinical T1D during the follow-up (PT1D, n=34), or (2) seroconverted to at least one islet autoantibody but were not diagnosed with T1D during the follow-up (P1Ab, n=27), and (3) children who remained autoantibody negative during the follow-up (CTRL, n=10) (**Table 1**). The lipidomics dataset comprised 153 lipid species. Identification of sources of variation in the PBMC lipidome dataset was performed using linear regression modeling, where the concentrations of lipids were regressed with various clinical variables such as age, gender and disease conditions. This analysis showed that the age of an individual indeed had a confounding effect (> 10% of explained variation, EV) on the lipidome **(Supplementary Figure S1)**. The effect of sex was, however, minimal (< 1% EV) **(Supplementary Figure S1)**.

**Table 1.** Demographic and clinical characteristics of the study population.

|  | **CTRL** | **P<sub>1</sub>Ab** | **PT<sub>1</sub>D** |
| --- | --- | --- | --- |
| **Number of subjects** | 10 | 27 | 34 |
| **Sex (Male, Female)** | (6, 4) | (16, 11) | (10, 24) |
| **Age at time of diagnosis (months, median ± SD)** | | | 53.0 ± 30.48 |
| **Age of first seroconversion (months, median ± SD)** | | 24.0 ± 20.12 | 14.0 ± 6.13 |
| **HLA risk\*** |  |  |  |
| **High risk** | 3 | 3 | 6 |
| **Moderate risk** | 1 | 14 | 17 |
| **Low or neutral risk** | 6 | 10 | 11 |

\*Risk groups defined in reference (Ilonen et al., 2016).
Abbreviations: children who progressed to T<sub>1</sub>D (PT<sub>1</sub>D); children who developed at least a single islet antibody but did not develop T<sub>1</sub>D during the follow-up (P<sub>1</sub>Ab); control (CTRL) subjects, who remained islet autoantibody negative during the follow-up; human leukocyte antigen (HLA); standard deviation (SD).

The PBMC lipidome was further explored using Sparse Partial Least Square Discriminant Analysis (sPLS-DA) (Le Cao et al., 2011), followed by univariate analysis (two sample t-test). The results suggest that lipid levels in PBMCs from the P1Ab and PT1D groups are different from CTRLs (**Figure 1A)**. Many classes of lipids, including cholesterol esters (CEs), lysophosphatidylcholines (LPCs), phosphatidylcholines (PCs), phosphatidylethanolamines (PEs), phosphatidylinositols (PIs), sphingomyelins (SMs), ceramides (Cers) and triacylglycerols (TGs) were significantly altered (AUC ~ 0.65, regression coefficient RC (>± 0.05) and VIP scores (Farrés et al., 2015) > 1 and T-test, p-value < 0.05) (**Figure 1A)**.

**Figure 1.**
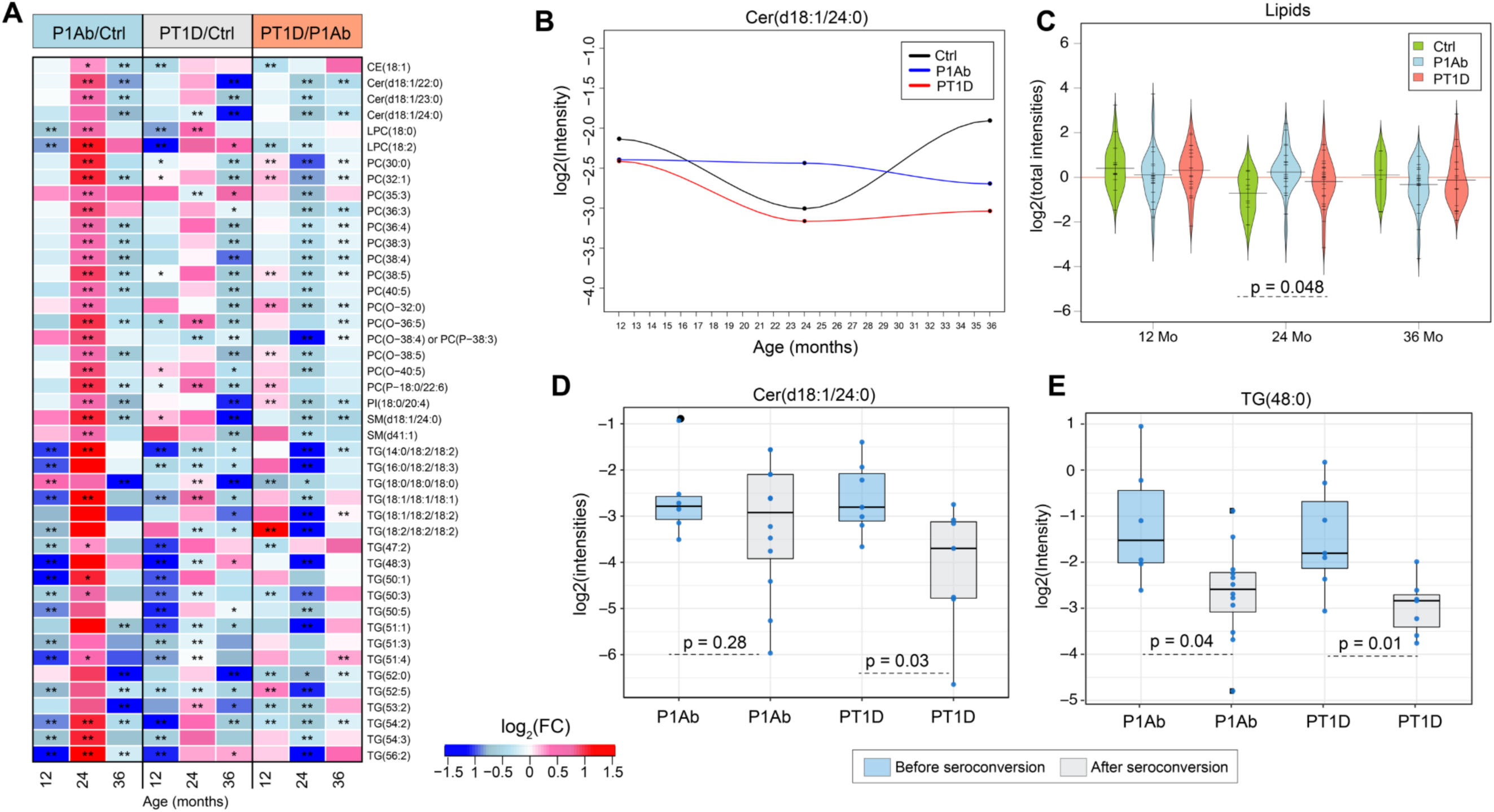
Lipid profiles in PBMCs during the follow-up. (A) Log2 fold changes (FCs) of lipid levels in P1Ab *vs*. CTRL, PT1D *vs*. CTRL and PT1D *vs*. P1Ab at 12, 24 and 36-months of age. ‘**’ denotes significant changes in lipid levels between the groups, as evaluated by univariate and multivariate analysis; ‘*’ denotes significant changes in lipid levels between groups, as evaluated by univariate analysis only. Red, blue and white color spectrum signifies up-, down-regulation and no change, respectively. (B) Profile of Cer(d18:1/24:0), which was significantly downregulated in PT1D. (C) Intensities of total lipids in CTRL, P1Ab and PT1D groups at 12, 24 and 36 months of age. The red, dotted line denotes the mean of the population. The black dashes in the bean plots represent individual subjects and their corresponding lipid levels, and the extended, black line denotes the group mean. (D-E) Boxplots showing the levels of Cer(d18:1/24:0) and TG(48:0) in subjects (marked with blue dots), before and after the seroconversion.

At 12 months of age, *i.e*. before the median age of seroconversion, the levels of some TG, PC, LPC and Cer species were lower in the PBMCs of the P1Ab and PT1D groups, as compared to CTRLs (**Figure 1A)**. This pattern was also observed at 36 months of age. However, at 24 months of age, there was a subtle increase or no change in these same lipid levels in the P1Ab and PT1D groups (**Figure 1A, Figure 1C, Supplementary Figure S2)**. This effect is most prominently seen in the P1Ab group, where the total lipid level was significantly higher (p=0.048) than in CTRL **(Figure 1C)**. Interestingly, this accumulation was transient, as the levels of these lipids returned to their previous (12-month timepoint) levels by 36 months (**Figure 1A, Figure 1C).** However, Cer(d18:1/24:0) was persistently decreased in the PT1D group as compared to P1Ab and CTRL groups (**Figure 1B)**. The time course profiles of selected lipids are shown in **Supplementary Figure S2.**

### PBMC lipidome before and after the first appearance of islet autoantibodies

Here we aimed to identify the molecular lipids which significantly changed between (1) prior to and (2) following on from seroconversion to islet autoimmunity in both the P1Ab and PT1D groups **(Figure 1D)**. Cer(d18:1/24:0), PC(36:3) and TGs with low carbon number and double bond count were downregulated in PT1D after seroconversion **(Supplementary Table S1)**. Total lipids in the P1Ab group (p = 5.581 × 10^−5^) and PT1D group (p = 9.803 × 10^−6^) were decreased after seroconversion (**Supplementary Figure S3)**.

### Polar metabolites of immune cells in progression to islet autoimmunity and T1D

We analyzed polar metabolites from PBMCs from the same samples as in the lipidomic analyses. By combining sPLS-DA and univariate analysis as above, we identified 25 polar metabolites which changed significantly (AUC ~ 0.60, RC (>± 0.05), VIP scores > 1 and T-test p-value < 0.05) (**Figure 2A)** between these groups at 12, 24, and 36 months of age. These metabolites can be divided into major chemical classes including carboxylic acids, amino acids, sugar derivatives, hydroxyacids, phenolic compounds, fatty acids and phosphate derivatives. Among the significantly-changing metabolites, comparative time course profiles of alanine and glutamic acids are shown in **Figures 2B** and **2D**.

**Figure 2.**
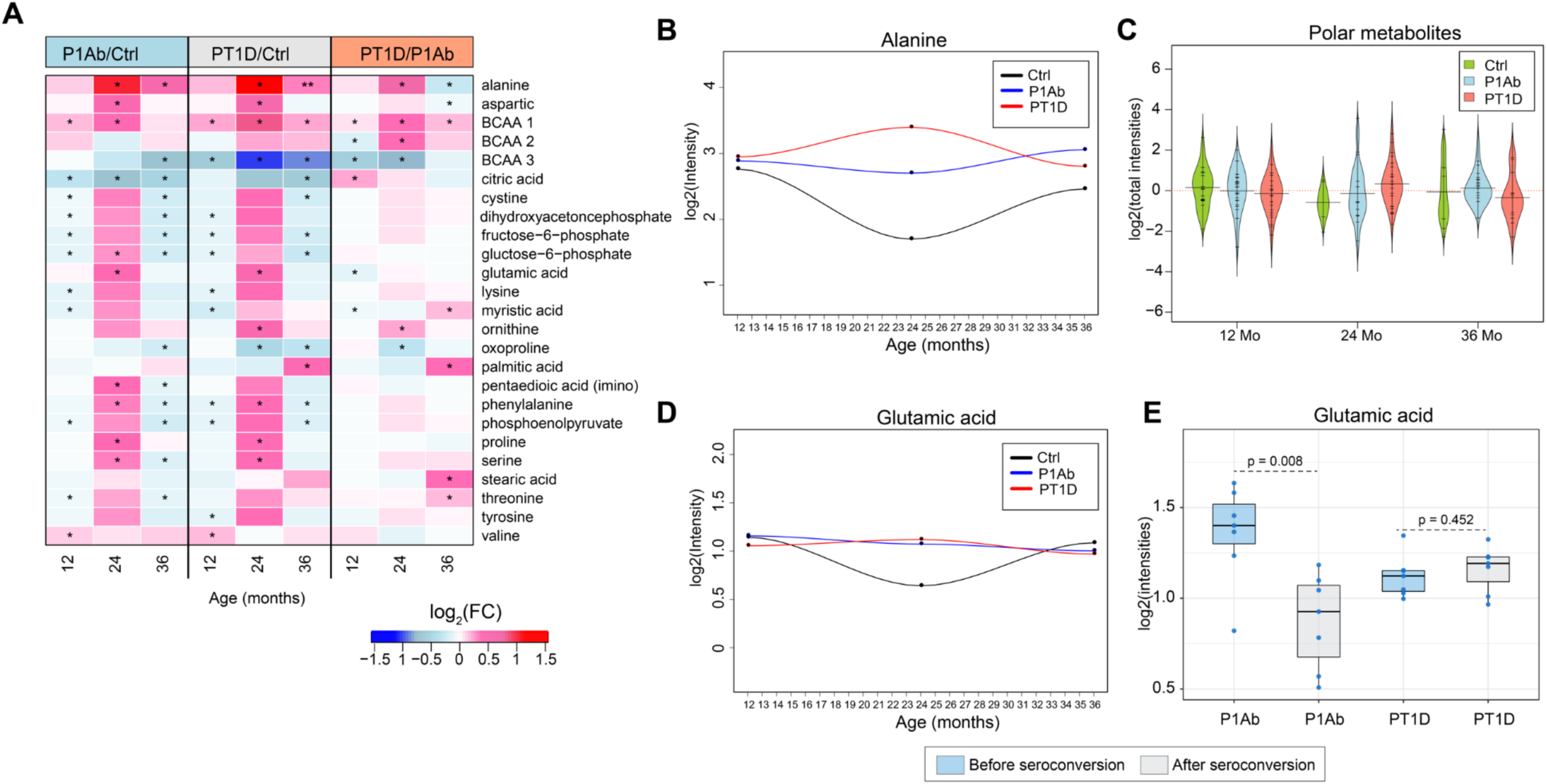
Polar metabolite profiles in PBMCs during the follow-up. (A) Log2 FCs in the levels of the polar metabolites in P1Ab *vs*. CTRL, PT1D *vs*. CTRL and PT1D *vs*. P1Ab at 12, 24 and 36-months of the follow-up. **’ denotes a significant change in the metabolite intensities between the groups, as evaluated by the univariate and multivariate analysis; ‘*’ denotes significant change in the metabolite intensities between the groups, as evaluated by univariate analysis only. Red, blue and white colors signify up-, down-regulation and no change respectively. (B) Time-course profile of alanine in CTRL, P1Ab and PT1D during the follow-up. (C) Log2 intensities of total polar metabolites as measured in CTRL, P1Ab and PT1D groups at 12, 24 and 36 months of age. The red, dotted line denotes the mean of the population. The black dashes in the bean plots represent individual subjects and their corresponding levels of total metabolites. The extended black line denotes the group mean. (D) Time-course profile of glutamic acid in CTRL, P1Ab and PT1D during the follow-up. (E) Levels of glutamic acid in each subject (marked with blue dots), before and after seroconversion.

At 12 months of age, a majority of the polar metabolites were downregulated in the P1Ab and PT1D groups as compared to the CTRL group. At 24 months there was a subtle increase in the levels of several amino acids including alanine, phenylalanine, proline, serine, threonine, cystine, lysine, glutamic and aspartic acid in the P1Ab and PT1D groups (**Figures 2A** and **2C**).

At 36 months of age, *i.e.*, after seroconversion in most children in the PT1D and P1Ab groups, a significant increase in several saturated fatty acid (FA) levels including stearic, myristic and palmitic acids was observed in the PT1D group *versus* the P1Ab group **(Figure 2A)**. When comparing metabolite levels before and after seroconversion (**Supplementary Figure S4**), glutamic acid was found to be significantly (p = 0.008) decreased after seroconversion in the P1Ab group (**Figure 2E**).

### Metabolic associations between immune cells and circulating metabolome

Next, we set out to examine how the metabolite profiles of PBMCs associated with their corresponding plasma profiles. We performed correlation analysis between the metabolites that were significantly altered in the PBMCs across the three study groups (**Figures 1** and **2)**, with their corresponding plasma levels, as reported previously (Lamichhane et al., 2018a; Lamichhane et al., 2018b). These two previous studies comprise an expanded group of individuals, a subsample of which was included in the present study.

In the CTRL group, at 12 months of age, the levels of PCs, LPCs, SMs and TGs in PBMCs were positively correlated (Spearman’s correlation coefficient, ρ > 0.70, p < 0.05) with their corresponding plasma levels (**Figure 3A)**. On the other hand, the polar metabolites were mostly inversely correlated, except alanine, glutamic acid and valine. At this same age, we found a distinct pattern of PC and TG levels in the PT1D group. Here, levels of cellular PCs were positively correlated to their corresponding plasma levels as in the CTRL group, whilst these PCs were negatively correlated with specific plasma TG levels (those of low carbon number and double bond count) (**Figure 3B)**. Likewise, the levels of plasma PCs were inversely correlated with cellular TG levels. In the P1Ab group, the association between cellular and plasma metabolites exhibited a different pattern, with the levels of cellular PCs being inversely correlated with their corresponding PCs and TG levels in the plasma **(Supplementary Figure S5)**.

**Figure 3.**
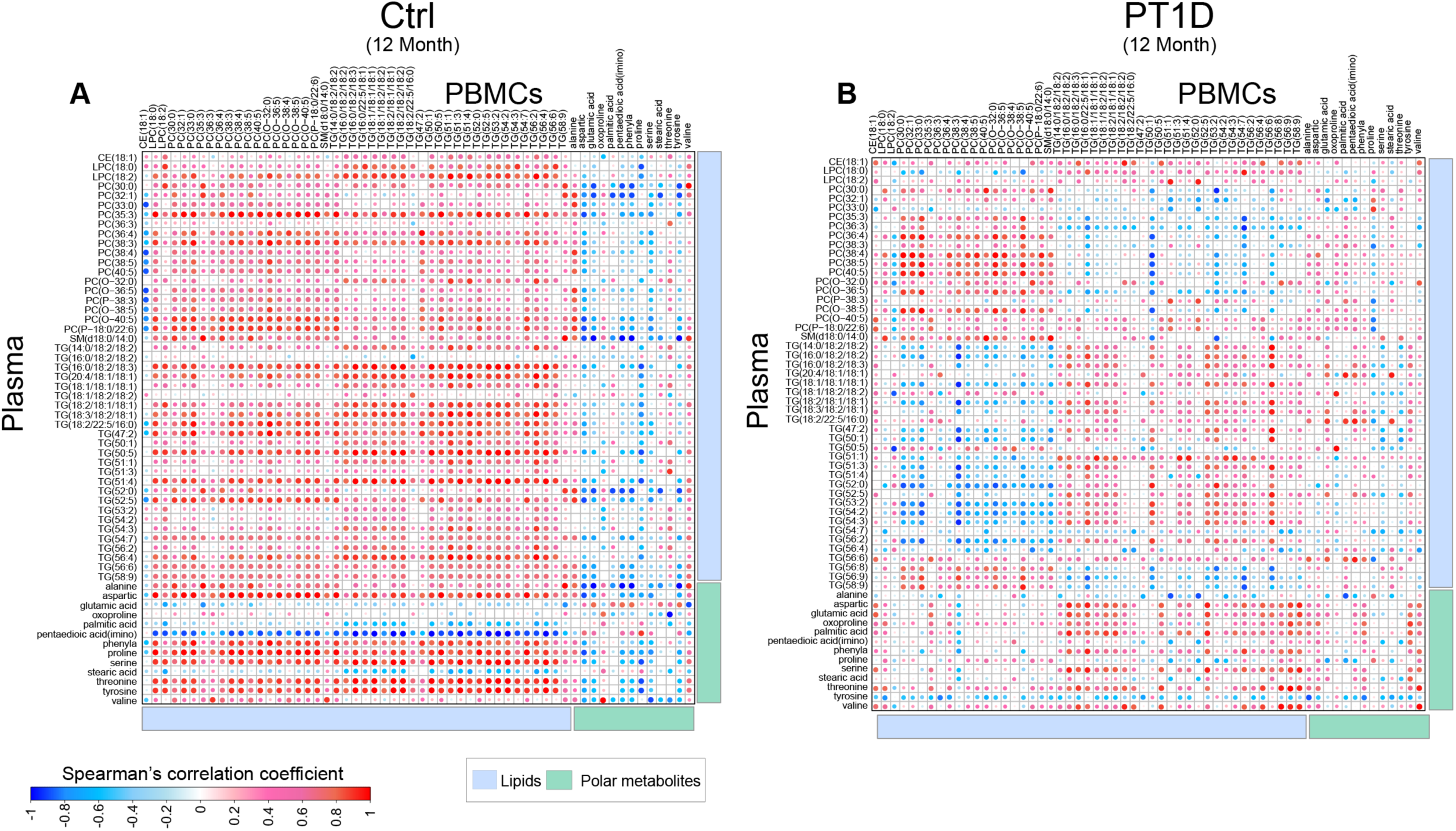
Spearman correlation between plasma and cellular metabolite levels in CTRL and PT1D groups at 12 months of age. Red, blue and white colors suggest positive, inverse and no correlation respectively.

However, in contrast, at the age of 36 months, the lipid profiles of PBMCs were predominantly positively correlated with their corresponding plasma in both the PT1D and P1Ab groups (**Supplementary Figures S5** and **S6)**. However, these associations were markedly different in the CTRL group, where predominantly inverse correlations between cellular and plasma lipids were observed, except in the case of TGs **(Supplementary Figure S6)**.

### Overrepresentation of metabolic pathways in progression to T1D

The lipids and polar metabolites of PBMCs which differed significantly between the three study groups were mapped against reference human metabolic pathways. The overrepresented metabolic subsystems/processes (for, *e.g.*, glycerolipid metabolism, pyruvate metabolism etc.) were selected based on a false discovery rate (FDR) < 0.05. Pathway impact score (PIS) for each subsystem was estimated (**Figure 4)**.

**Figure 4.**
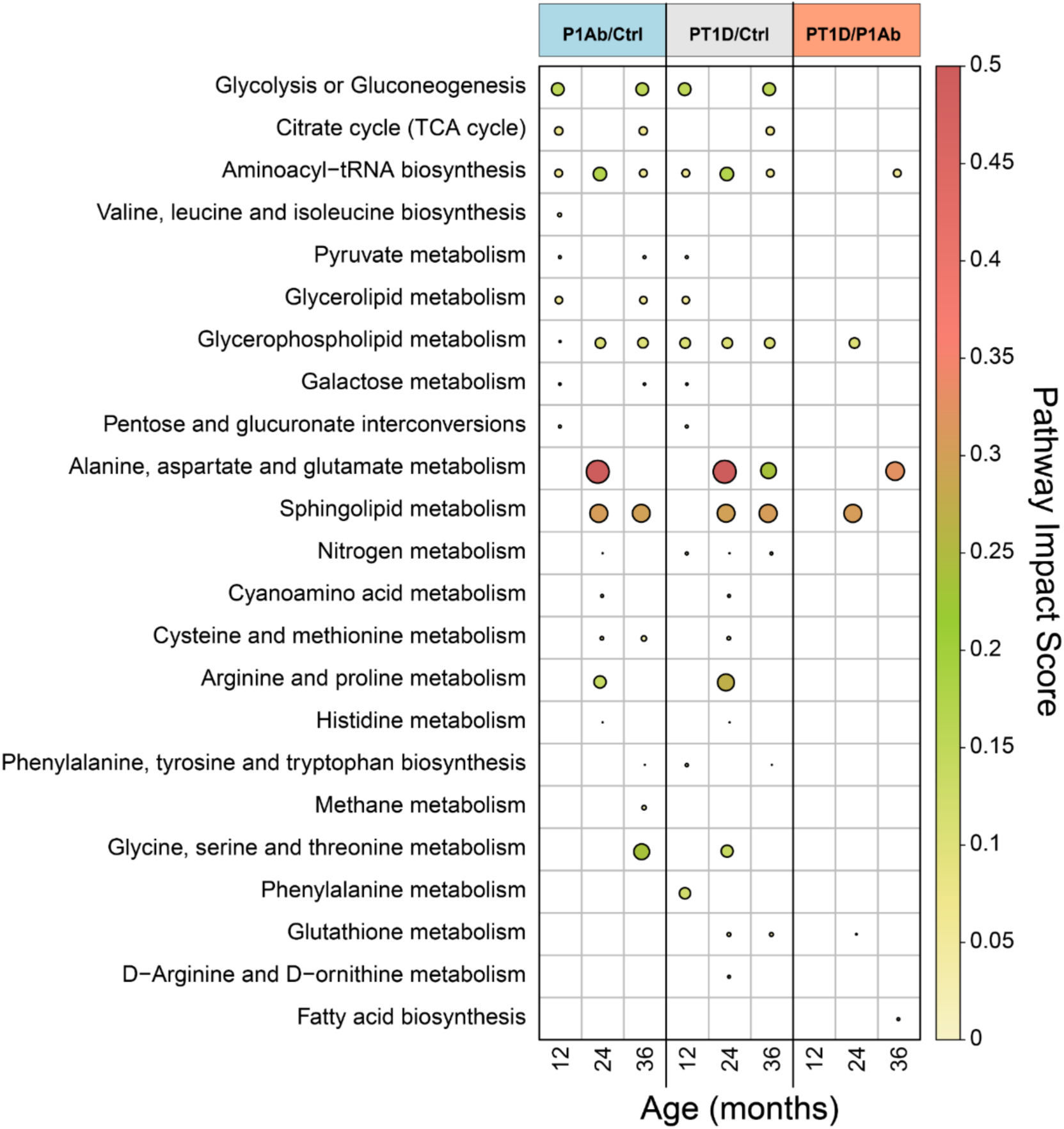
Overrepresentation analysis of metabolic pathways in PBMCs in CTRL, P1Ab and PT1D groups. The plot shows the significant pathway impact scores (PIS) (FDR < 0.05) of each metabolic subsystem/pathway during the follow-up. Red, green and yellow colors denote high, intermediate and low impacts, respectively.

Various core metabolic processes were overrepresented in the immune cells in the PT1D and P1Ab groups at 12 months of age, *i.e*, preceding islet seroconversion and overt T1D. These include central carbon metabolism (CCM), sugar metabolism, amino-acid biosynthesis (valine, leucine and isoleucine) and glycerophospholipid (GPL) metabolism (**Figure 4)**. Other metabolic subsystems such as alanine, aspartate and glutamate metabolism (AAG) (PIS~0.4), sphingolipid metabolism (SMM) (PIS~0.4), GPL metabolism (PIS~0.15), arginine and proline metabolism (PIS~0.25), and aminoacyl-tRNA biosynthesis (PIS~0.2) were overrepresented in the P1Ab and PT1D groups at 24 months of age. Moreover, SMM, GPL, AAG, CCM, glycine, serine and threonine (GST) metabolism were especially overrepresented at 36 months of age, among which AGG, SMM, GPL and FA biosynthesis were overrepresented in the PT1D group as compared to the P1Ab group (**Figure 4)**.

### Metabolic modeling of sphingolipid metabolism in islet autoimmunity and T1D

Given that: (1) in our previous study we observed persistent downregulation of plasma sphingolipids in children who progressed to T1D (Lamichhane et al., 2018a; Oresic et al., 2008), (2) SMM was overrepresented in PT1D PBMCs in the present study, and (3) we recently found that prenatal chemical exposure modulates postnatal sphingomyelin levels and increases T1D risk (McGlinchey et al., 2019), we therefore further examined PBMC SMM using genome-scale metabolic modeling **(Supplementary Figure S7)**. The objective of metabolic modeling, in this case, was to identify the key regulators within SMM in progression to T1D. This extended analysis was performed by integrating publicly-available transcriptomics datasets obtained from the two related cohorts and the metabolomics dataset from the present study (see **Methods**). By using metabolomics dataset from the present study, we devised a confidence score for each metabolite as being either present or absent in the PBMC metabolic model. The constraints for model exchange/input reactions were derived using the metabolomics data from the present study.

Reporter metabolite (RM) analysis showed that glucosyl-, lactosyl- and galactosylceramides were significantly up-regulated in the PT1D group as compared to P1Ab (**Figure 5B)** and CTRL **(Supplementary Figure S8)**. These changes were, however, not observed when comparing the P1Ab group against CTRLs. Instead, leukotrienes LTB4, HETE and EpOME derivatives (markers of inflammation) were significantly up-regulated **(Supplementary Figure S9)**. This modeling of SMM suggests that, in the PT1D group, cellular ceramides are converted to glycoceramides, thus decreasing the free ceramides in the cells.

**Figure 5.**
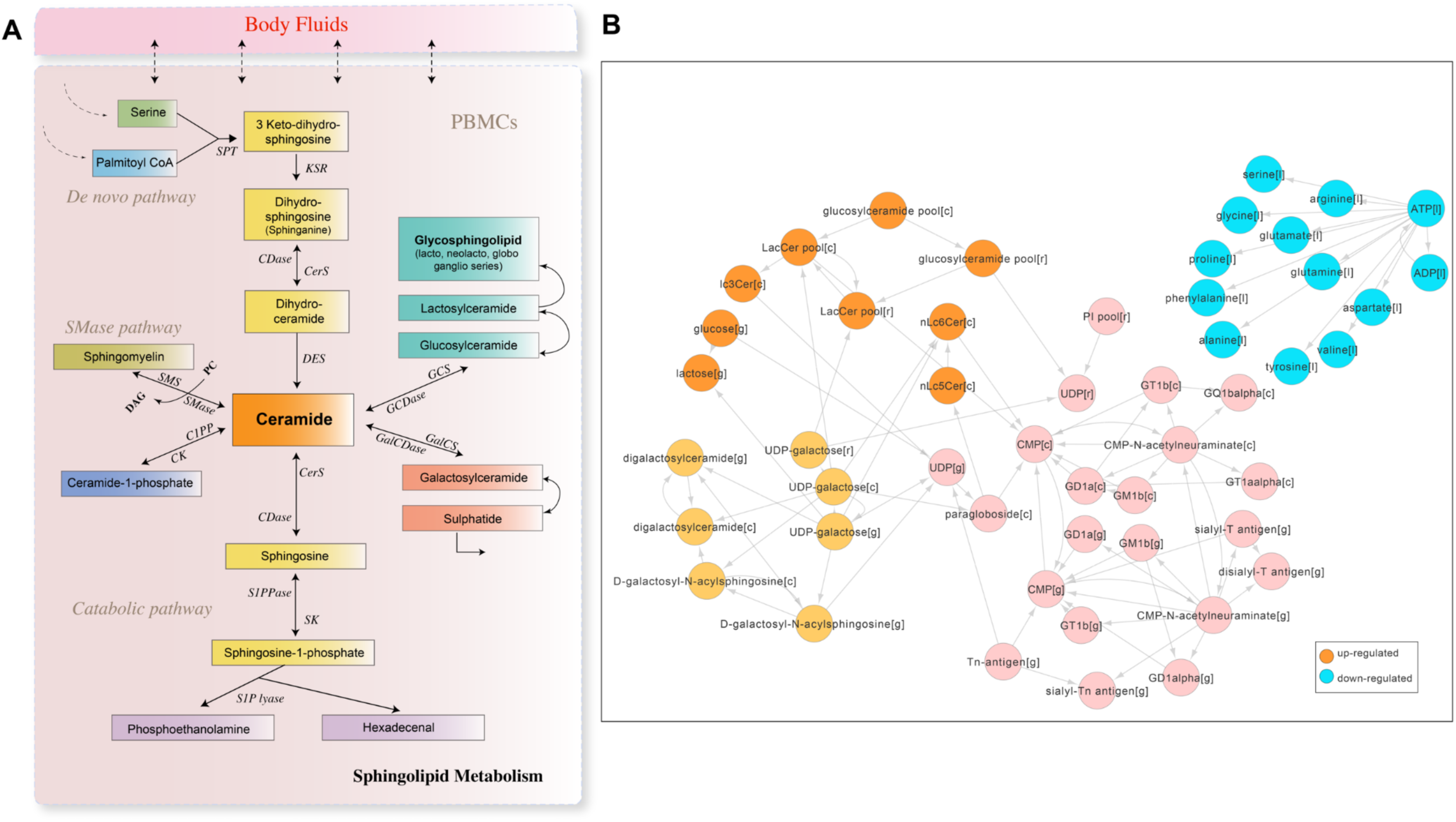
Regulation of sphingomyelin pathways in progression to islet autoimmunity and overt T1D. (A) Canonical pathways of sphingolipid metabolism in human. (B) Reporter metabolites predicted for PBMCs that changed significantly (FDR < 0.05) between PT1D and P1Ab. The orange and cyan colors denote up- and down-regulation of the reporter metabolites respectively.

In order to confirm these findings, we analyzed six glycoceramides from the lipidomics dataset: HexCer(d18:1/16:0), HexCer(d18:1/22:0), HexCer(d18:1/24:0), LacCer(d18:1/12:0), LacCer(d18:1/14:0), LacCer(d18:1/16:0). At 36 months of age, HexCer(d18:1/16:0) and HexCer(d18:1/22:0) were found upregulated in PT1D as compared to P1Ab, with a similar trend observed for HexCer(d18:1/24:0) and LacCer(d18:1/12:0), LacCer(d18:1/14:0), while LacCer(d18:1/16:0) displayed an opposite trend (**Figure 6)**. There was congruence between the predicted RMs (**Figure 5B)** and the glycoceramide levels measured in PT1D at 36 months of age. At this age, the ceramides (*i.e.*, Cer(d18:1/24:0) and Cer(d18:1/22:0)) were down-regulated in PT1D as compared to P1Ab (**Figure 1A)**.

**Figure 6.**
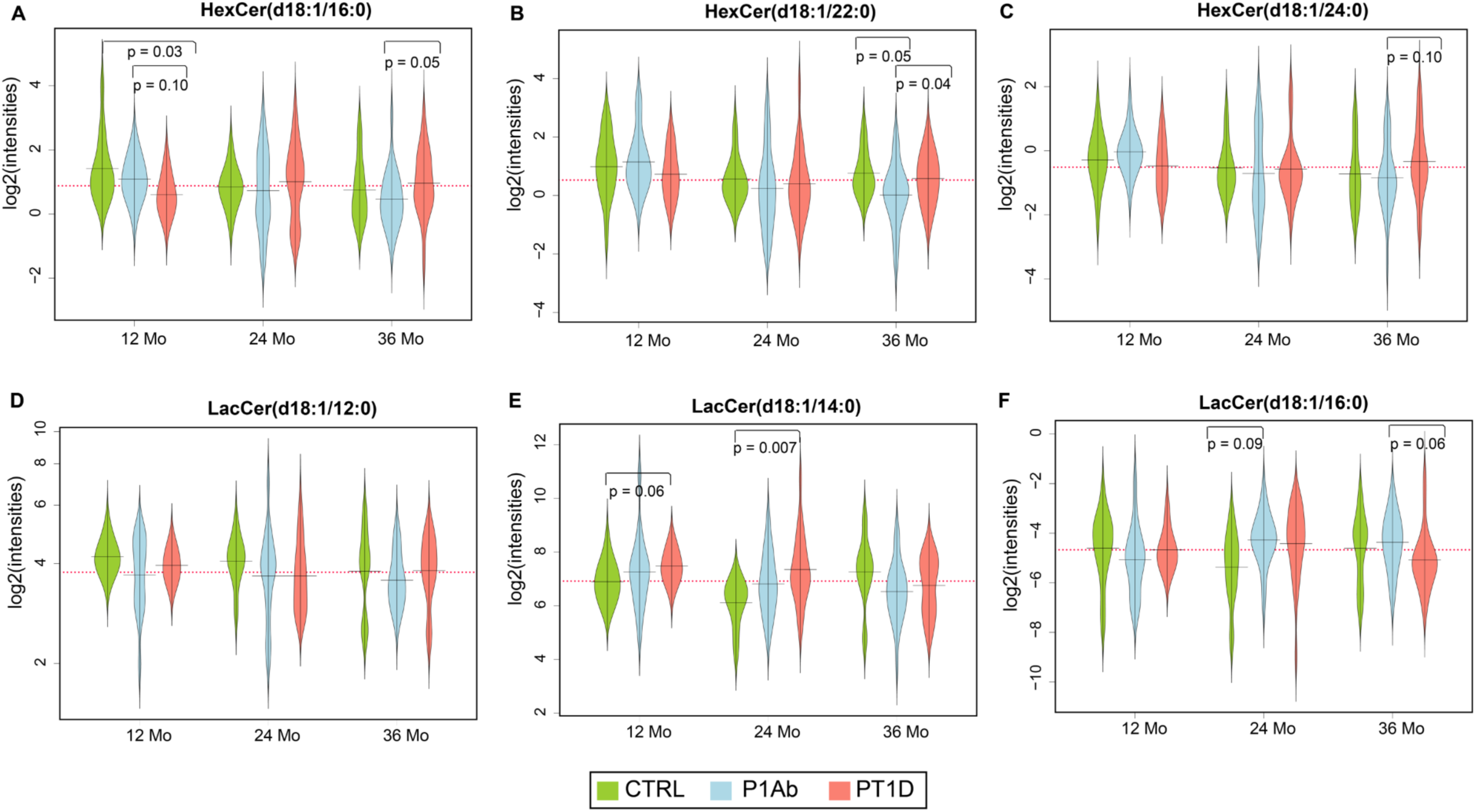
Beanplots showing the levels of hexosyl- and lactosylceramides in PBMCs. A-F) Levels of glycoceramides in CTRL, P1Ab and PT1D at 12,24 and 36 months (Mo). The red, dotted line denotes the mean of the population. The black dashes in the bean plots represent the group mean. ‘p’ represents the p-values obtained from a t-test. p-values < 0.1 are marked.

In order to understand the significance of glucosyl-, lactosyl- and digalactosylceramide production, we optimized two, key cytoplasmic reactions in SMM: (1) production of glucosylceramides from ceramides and D-glucose **(Supplementary Figure S10A)**, (2) formation of digalactosylceramide from N-acyl-sphingosine and D-galactose in the three groups, and thereby recorded the flux changes across the different metabolic subsystems/pathways **(Supplementary Figure S10B).** The results suggest that, among several affected metabolic processes, mitochondrial and endoplasmic reticular transport of substrates might be essential for glycoceramide production. In addition, marked changes in GPL, glycolysis/gluconeogenesis and amino acid metabolism were observed.

## DISCUSSION

We identified differences in the metabolomes of PBMCs isolated from (1) T1D progressors (PT1D), (2)non-progressors who converted to one or more islet autoantibodies during the follow-up (P1Ab) and (3) controls (CTRL) that remained autoantibody negative throughout follow-up. In line with previous findings in plasma (La Torre et al., 2013; Lamichhane et al., 2018a; Oresic et al., 2013; Oresic et al., 2008), these differences were observed even before the first appearance of islet autoantibodies.

The observed metabolic changes in PBMCs were dominated by glycerophospholipids, triacylglycerols, sphingomyelins, ceramides, amino acids and fatty acids. During the first year of life, *i.e.*, before the median age of seroconversion, a majority of lipids and polar metabolites measured in PBMCs were down-regulated in the P1Ab and PT1D groups as compared to CTRL. The occurrence of low levels of TGs of low double bond count and carbon number (*i.e.*, TGs enriched with saturated fatty acids such as palmitate and myristate) suggest an impairment of *de novo* lipogenesis in PBMCs as being potentially involved in progression towards T1D. Negative associations between PC and TG levels in (1) PBMCs and (2) blood plasma of children who progressed to T1D suggest that in PT1D, the levels of PCs and TGs in the immune cells might depend on their corresponding plasma levels or *vice versa*.

At 24 months, *i.e.*, a period coinciding with seroconversion in a majority of children in the PT1D and P1Ab groups, there was an increase particularly in the levels of lipids and amino acids as compared to the control group. There is evidence that amino acids, in particular branched chain amino acids (BCAAs), contribute to dysregulated lipid metabolism (White et al., 2018). We observed higher levels of BCAAs and related metabolites, including glutamic acid and alanine, in circulating PBMCs during this period, which is indicative of compromised amino acid catabolism and lipogenesis (Green et al., 2016). In parallel, elevated BCAAs also promote pro-inflammatory signaling in PBMCs (Zhenyukh et al., 2017). Increased concentrations of BCAAs in PBMCs were demonstrated to coincide with the period of seroconversion in the PT1D and P1Ab groups. This observed transient phenomenon suggests that increased BCAAs contribute to immune dysfunction in children who later progress to T1D.

There is growing evidence of abnormal SMM in progression to and in early T1D (Galadari et al., 2013; Greenhill, 2018; Holm et al., 2018). Differential analysis of PBMC metabolites in our study identified changes in the levels of ceramides and sphingomyelin species in the PT1D group as compared to CTRL. Sphingomyelin cleavage and ceramide synthesis is one of the important mechanisms involved in the regulation of immune cell function (Beyersdorf and Muller, 2015). Serine and palmitic acid are precursors for *de novo* sphingomyelin biosynthesis (Gault et al., 2010). We found that serine concentrations were decreased in T1D progressors after seroconversion. Previous studies by us and others have also reported decreased plasma levels of SMs in children who later progressed to T1D as well as in children with newly-diagnosed T1D (Lamichhane et al., 2018a; Orešic et al., 2008; Sorensen et al., 2010). Our pathway overrepresentation analysis also corroborates that SMM is overrepresented in the P1Ab and PT1D groups. This suggests that altered sphingolipid metabolism in immune cells, as well as in circulation, is a hallmark of progression to overt T1D.

Genome-scale metabolic modeling predicted that ceramide pathways leading to synthesis of glucosyl-, lactosyl- and galactosylceramides were upregulated in PT1D as compared to P1Ab. This prediction was confirmed by way of the observed increased levels of these glycosphingolipids in the PT1D group. This strongly implies that ceramides, found down-regulated in the PT1D group of our study, are converted to glycoceramides after seroconversion in children who later progress to T1D. In agreement with this, we found that the gene expression of glucosylceramide synthase (EC 2.4.1.80), a rate limiting enzyme in the conversion of ceramides to glucosylceramide and downstream glycosphingolipids, was also increased. Glycoceramides, particularly glucosylceramide, play an important role in the control of immune responses (Brennan et al., 2014). Previous studies have shown that these glycosphingolipids modulate β-cell immune receptor signaling (Boslem et al., 2012; Osterbye et al., 2010) as well as aggravate systemic inflammation responses (Mobarak et al., 2018; Pandey et al., 2017). Elevated glycoceramide levels in PBMCs in autoantibody-positive children who later progress to T1D, as observed in our study, therefore point to a specific sphingolipid pathway in immune cells which contributes to T1D pathogenesis.

Taken together, our results suggest that progression to T1D is accompanied by metabolic abnormalities in PBMCs. These changes may be related to impaired *de novo* lipogenesis, amino acid metabolism, GPL metabolism and SMM. Since specific differences were also observed between progressors and non-progressors to T1D after their seroconversion to islet autoimmunity, our findings also highlight specific pathways in immune cells, such as sphingolipid metabolism, which appear to play a role in protection from and progression to T1D. It remains to be established how these and other pathways are altered in specific immune cell subtypes.

## METHODS

### Study design and protocol

In this study, the samples were obtained from the Finnish Type 1 Diabetes Prevention and Prediction Study (DIPP) (Kupila et al., 2001). The DIPP study has screened more than 230,000 newborn infants for HLA-conferred susceptibility to T1D in three university hospitals; those at Turku, Tampere, and Oulu in Finland (Haller and Schatz, 2016). The subjects involved in the current study were chosen from the subset of DIPP children which were from the city of Tampere, Finland. The study protocol was approved by the ethics and research committee of the participating university and hospital. The study was conducted according to the guidelines in the Declaration of Helsinki. All families provided written, informed consent for participation in the study. Here, longitudinal samples for each child were collected between 1998 and 2012. For each child, longitudinal samples for PBMC metabolomic analysis were obtained at 12, 24 and 36 months of age.

This study comprises samples (n=137 for lipidomics and n=134 for polar metabolites) from 71 children, divided into three groups, being: (1) 27 children who seroconverted to at least one islet autoantibody but were not diagnosed with T1D during the follow-up (P1Ab), (2) 34 children seroconverted to more than one islet autoantibody and subsequently developed T1D (PT1D), and (3) 10 controls (CTRL), *i.e.*, children who remained islet autoantibody negative during follow-up. The three study groups were matched for HLA-associated risk for T1D, sex and age. Selected characteristics of the subjects involved in this study are listed in **Table 1**. In this study, non-fasting blood samples were collected, plasma was prepared within 3 hours of sample collection and stored at −80°C until analyzed. PBMCs were purified using BD Vacutainer® CPT™ Cell Preparation Tube with sodium citrate (Becton, Dickinson and Company, Franklin Lakes, NJ) according to manufacturer’s instructions. Collected whole blood was let to cool down for 15-20 minutes in the blood collection tube at room temperature. The blood samples were centrifuged at 2800 rpm for 20 minutes, after which the layer of mononuclear cells was suspended into the plasma and the suspension was transferred to a new tube. The sample was centrifuged again at 1700 rpm for 15 minutes. The samples were then divided into cryotubes and snap frozen in liquid nitrogen. The tubes were kept in liquid nitrogen overnight and then stored at −70 °C. For the metabolomic analysis, the samples were thawed on ice and the initial lysing procedures performed in a cold room to prevent changes to the metabolites from the cells or proteins. The cell pellets were resuspended in ice cold saline (0.9% NaCl) by gently pipetting up and down. 25 µL of cells were then aliquoted for metabolomic analysis and stored at −80°C. The total protein content in cells was measured by the Bradford method (Bradford, 1976).

### HLA genotyping

Screening for HLA-conferred susceptibility to T1D was performed using cord blood samples. The HLA-genotyping was performed using a time-resolved, fluorometry-based assay for four alleles using lanthanide chelate-labeled, sequence-specific oligonucleotide probes detecting DQB1*02, DQB1*03:01, DQB1*03:02, and DQB1*06:02/3 alleles (Ilonen et al., 1996). The carriers of genotypes DQB1*02/DQB1*03:02 or DQB1*03:02/x genotypes (here x = DQB1*03:01, DQB1*06:02, or DQB1*06:03 alleles) were categorized as being eligible and recruited for the DIPP follow-up program in Tampere until 3 years of age.

A more extensive HLA genotyping was performed for the children participating this study. This genotyping defined all common European HLA-DR-DQ haplotypes at low resolution and at higher resolution haplotypes where this was relevant for estimation of the risk for T1D conferred, e.g. HLA-DR4 subtypes in DR4-DQ8 haplotypes. In a series of 2,991 family trios from the Finnish Pediatric Diabetes Register, the genotype risks were defined and genotypes were combined into 6 groups from (strongly protective) to 5 (high risk) which did not overlap for 95% confidence intervals of their OR values for T1D (Ilonen et al., 2016).

### Detection of islet autoantibodies

The children with HLA-conferred genetic susceptibility were prospectively observed for levels of T1D associated autoantibodies (ICA, IAA, IA-2A, and GADA). These autoantibodies were assayed from plasma samples taken at each follow-up visit as previously described (Siljander et al., 2009). ICA levels were determined using an approved, immunofluorescence assay with a detection limit of 2.5 Juvenile Diabetes Foundation Units (JDFU) (Bottazzo et al., 1974). GADA and IAA levels were quantified using specific radiobinding assays, the threshold of positivity being 5.36 and 3.48 relative units (RU) respectively (Ronkainen et al., 2001; Savola et al., 1998b). Similarly, IA-2A levels were measured with a radio-binding assay with a threshold of 0.43 RU (Savola et al., 1998a).

### Analysis of molecular lipids

The samples were randomized and extracted using a modified version of the previously-published Folch procedure (Folch et al., 1957). Briefly, 150 µL of 0.9% NaCl was added to the cell pellets, and samples were then vortex mixed and ultrasonicated for 3 minutes. Next, 20 µL of the cell suspension was mixed with 150 µL of the 2.5 µg mL^−1^ internal standards solution in ice-cold CHCl3:MeOH (2:1, v/v). The internal standard solution contained the following compounds: 1,2-diheptadecanoyl-sn-glycero-3-phosphoethanolamine (PE (17:0/17:0)), N-heptadecanoyl-D-erythro-sphingosylphosphorylcholine (SM(d18:1/17:0)), N-heptadecanoyl-D-erythro-sphingosine (Cer(d18:1/17:0)), 1,2-diheptadeca-noyl-sn-glycero-3-phosphocholine (PC(17:0/17:0)), 1-heptadecanoyl-2-hydroxy-sn-glycero-3-phosphocholine (LPC(17:0)) and 1-palmitoyl-d31-2-oleoyl-sn-glycero-3-phosphocholine (PC(16:0/d31/18:1)). These were purchased from Avanti Polar Lipids, Inc. (Alabaster, AL, USA). In addition, triheptadecanoin (TG(17:0/17:0/17:0)) was purchased from (Larodan AB, (Solna, Sweden). The samples were vortex mixed and incubated on ice for 30 min after which they were centrifuged at 7800 × g for 5 min. Finally, 60 µL from the lower layer of each sample was collected and mixed with 60 µL of ice-cold CHCl3:MeOH (2:1, *v/v*) in an LC vial.

The UHPLC-QTOFMS analyses were done in a similar manner to that described earlier, with some modifications (Nygren et al., 2011; Pedersen et al., 2018). The UHPLC-QTOFMS system was from Agilent Technologies (Santa Clara, CA, USA) combining a 1290 Infinity LC system and 6545 quadrupole time-of-flight mass spectrometer (QTOFMS), interfaced with a dual jet stream electrospray (dual ESI) ion source. MassHunter B.06.01 software (Agilent Technologies, Santa Clara, CA, USA) was used for all data acquisition and MZmine 2 was used for data processing (Pluskal et al., 2010).

Chromatographic separation was performed using an Acquity UPLC BEH C18 column (100 mm × 2.1 mm i.d., 1.7 µm particle size) and protected using a C18 precolumn, both from Waters Corporation (Wexford, Ireland). The mobile phases were water (phase A) and acetonitrile:2-propanol (1:1, *v/v*) (phase B), both containing 1% 1M ammonium acetate and 0.1% (*v/v*) formic acid ammonium acetate as ionization agents. The LC pump was programmed at a flow rate of 0.4 mL min^−1^ and the elution gradient was as follows: from min 0–2, the percentage of phase B was modified from 35% to 80%, from min 2-7, the percentage of phase B was modified from 80% to 100% and then, the final percentage was held for 7 min. A post-time of 7 min was used to regain the initial conditions for the next analysis. Thus, the total analysis time per sample was 21 min (including postprocessing). The settings of the dual ESI ionization source were as follows: capillary voltage 3.6 kV, nozzle voltage 1500 V, N_2_ pressure in the nebulizer 21 psi, N_2_ flow rate and temperature as sheath gas 11 L min^−1^ and 379 °C, respectively. Accurate mass spectra in the MS scan were acquired in the m/z range 100–1700 in positive ion mode.

Identification of lipids was carried out by combining MS (and retention time), MS/MS information, and a search of the LIPID MAPS spectral database (http://www.lipidmaps.org). MS/MS data were acquired in both negative and positive ion modes in order to maximize identification coverage. Additionally, some lipids were verified by injection of commercial standards. The confirmation of a lipid’s structure requires the identification of hydrocarbon chains bound to its polar moieties, and this was possible in some cases. This identification was carried out in pooled cell extracts, and with this information, an in-house database was created with m/z and retention time for each lipid. This in-house database was used for processing data by MZmine 2 (Pluskal et al., 2010). Glycoceramides were identified based on their accurate mass, MS/MS analysis and retention times. Further verification of some of the ceramides was possible with authentic standards. The peak areas were normalized with Cer(d18:1/17:0) and quantified against the calibration curve of Cer(d18:1/18:0).

The peak area obtained for each lipid was normalized with lipid-class-specific internal standards and with total content of protein. A (semi) quantitation was performed using lipid-class-specific calibration curves. Pooled cell extracts were used for quality control, in addition to in-house plasma. The raw variation of the peak areas of internal standards in the samples was, on average, 12.0%, and the RSD of retention times of identified lipids across all samples was on average 0.47%. The RSD of the concentrations of the identified lipids in QC samples and pooled extracts was on average 23%.

### Analysis of polar metabolites

To the remaining 75 µL aliquot, 225 µL of ice-cold MeOH (LC-grade, Honeywell) containing the following internal standards (all from Sigma Aldrich): Heptadecanoic acid (5 ppm), DL-valine-d8 (1 ppm), and succinic acid-d4 (1 ppm) was added. The samples were then sonicated in an ice bath for 30 s prior to centrifugation (5500 g, 5 min). 250 µL of the supernatant was transferred to a 2 mL glass autosampler vial. The pellet was stored at –20 °C for protein analysis. The protein content was measured by the Bradford method. The supernatant was dried under a stream of nitrogen at 45 °C. Prior to the mass spectrometry measurements, the samples were derivatized using a two-step procedure. Initially the samples were methoximated by incubating the samples with methoxyamine hydrochloride (25 µL, 20mg/mL in pyridine, Sigma Aldrich) at 45 °C for 1 h. MSTFA (25 µL, Sigma Aldrich) was then added and the samples were incubated for a further 1 h. A retention index standard containing straight chain, even alkanes (n 10-40, 10 µL, Sigma Aldrich) was added. The derivatized samples were analyzed using gas chromatography (Agilent 7890B) coupled to a single quad mass spectrometer (5977B). The metabolites were separated using a 30 m × 0.25 mm (ID) with a film thickness of 0.25 µm HP-5 (Agilent). A guard column (10 m) with an ID of 0.25 mm was used. 1 µL of the sample was injected in splitless mode with an inert glass liner (Agilent) held at a temperature of 240 °C. The GC was set to constant flow mode (1.2 mL/min) using helium (Aga) as the carrier gas. The GC oven was programed as follows: 50 °C (isothermal for 0.2 min), then 7 °C/min until 240 °C, then 20 °C/min until 300 °C (isothermal for 5 min). The transfer line was held at 260 °C for the whole run. The ion source was set to electron ionization mode and held at 230 °C and the quadrupole at 150 °C. Due to the large number of analytes quantified, the samples were injected twice. The first run quantified the amino acids and the second run quantified all other components. The MSD was set up in select ion monitoring mode to maximize sensitivity. The ions monitored can be found in **Supplementary Table S2**.

### Data preprocessing

Lipidomics data processing was performed using open source software MZmine 2.33 (Pluskal et al., 2010). The following steps were applied in the processing: 1) Crop filtering with a m/z range of 350 – 1700 m/z and a RT range of 2.0 to 12 min, 2) Mass detection with a noise level of 1200, 3) Chromatogram builder with a min time span of 0.08 min, min height of 1000 and a m/z tolerance of 0.006 m/z or 10.0 ppm, 4) Chromatogram deconvolution using the local minimum search algorithm with a 70% chromatographic threshold, 0.05 min minimum RT range, 5% minimum relative height, 1200 minimum absolute height, a minimum ratio of peak top/edge of 1 and a peak duration range of 0.08 - 5.0, 5) Isotopic peak grouper with a m/z tolerance of 5.0 ppm, RT tolerance of 0.05 min, maximum charge of 2 and with the most intense isotope set as the representative isotope, 6) Join aligner with a m/z tolerance of 0.008 or 10.0 ppm and a weight for of 2, a RT tolerance of 0.1 min and a weight of 1 and with no requirement of charge state or ID and no comparison of isotope pattern, 7) Peak list row filter with a minimum of 12 peaks in a row (= 10% of the samples), 8) Gap filling using the same RT and m/z range gap filler algorithm with an m/z tolerance of 0.006 m/z or 10.0 ppm, 9) Identification of lipids using a custom database search with an m/z tolerance of 0.006 m/z or 10.0 ppm and a RT tolerance of 0.1 min, 10) Normalization using lipid-class-specific internal standards and (semi) quantitation with lipid-class-specific calibration curves, 11) Normalization with total protein amount 12) Data imputation of missing values were done with half of the row’s minimum.

The GC-QMS data was processed in MassHunter Quant (v8, Agilent technologies) The peaks were manually checked and corrected if needed for correct integration. Quantification was performed using the ion listed **Supplementary Table S2**. Standard curves were used to quantify each metabolite using the assigned internal standards. Metabolites which had a CV greater than 30% in the pooled QC sample or fell below the limit of quantification were excluded from subsequent analysis.

### Statistical methods

The lipidomics and polar metabolites datasets were divided into three study groups: CTRL, P1Ab, and PT1D **(Table 1, Supplementary Figure S11)**. The age of the subject was calculated as a time difference between the date of the sample withdrawn and their date of birth. If more than two samples from the same case matched a time interval, the closest was selected. Each group was divided into three age groups of 12, 24 and 36 months **(Supplementary Figure S11)**. The data were log2 transformed. Homogeneity of the samples were assessed by principal component analysis (PCA) (Carey et al., 1975) and no outliers were detected. The log2-normalized intensities of the total identified lipids and polar metabolites in the subjects are shown in (**Supplementary Figures S12** and **S13)**.

The R statistical programming language (R Development Core Team, 2018) was used for data analysis and visualization. PCA was performed using *‘prcomp’* function included in the *‘stats’* package. The effect of different factors such as age, gender and conditions on the lipidomics dataset was evaluated. The data was centered to zero mean and unit variance. The relative contribution of each factor to the total variance in the dataset was estimated by fitting a linear regression model, where the normalized intensities of metabolites were regressed to the factor of interest, and thereby median marginal coefficients (R^2^) were estimated **(Supplementary Figure S1)**. This analysis was performed using *‘scater’* package.

Sparse Partial least squares Discriminant Analysis (sPLS-DA) (Le Cao et al., 2011) models comparing P1Ab *vs.* CTRL, PT1D *vs.* CTRL, and PT1D *vs.* P1Ab groups, paired at 12, 24 and 36 months were developed and Variable Importance in Projection (VIP) scores (Farrés et al., 2015) were estimated. sPLS-DA modeling was performed using the ‘*splsda’* function coded in the *‘mixOmics v6.3.2’* package. Moreover, the sPLS-DA models were cross-validated (Westerhuis et al., 2008) by 7-fold cross-validation and models diagnostics were generated using *‘perf’* function. The multivariate analysis was followed by univariate, Two-Sample t-testing using the *‘t.test’* function, to identify significant differences in the metabolite intensities between CTRL, P1Ab or PT1D groups. All metabolites that passed one or more criteria for variable selection, *i.e.*, with sPLS-DA model area under the ROC curve (AUC) >= 0.6; RC (>± 0.05), VIP scores > 1 or T-test; p-value < 0.05) were listed as significant. Spearman’s correlation coefficient (ρ) was calculated using the *‘rcorr’* function implemented in the *‘Hmisc’* package. p-values were subjected to False Discovery Rates (FDR) adjustment using ‘*p-adjust’*. Loess regression was performed using *‘loess’* function deployed in the ‘stats’ package. *‘Heatmap.2’, ‘boxplot’, ‘beanplot’*, ‘*gplot*’, and ‘*ggplot2*’ libraries/packages were used for data visualization.

Pathway overrepresentation analysis (POA) was performed using the MetaboAnalyst 4.0 web platform (Chong et al., 2018) using the ‘Pathway Analysis’ module. Those metabolites changing significantly between different groups were listed and mapped to the human metabolic network as a background; ‘relative-betweenness Centrality’ was selected for ‘pathway topology analysis’, and global hypergeometric test (GHT) was performed. GHT estimated the relative significance of the overrepresented pathways against the background KEGG pathways (Kanehisa et al., 2013) for *Homo sapiens*. The metabolic subsystems/pathways with FDR < 0.05 is reported in (**Figure 4)**. The Pathway Impact Scores (PIS) were estimated by the metabolomics pathway analysis (MetPA) tool (Xia and Wishart, 2010) encoded in MetaboAnalyst 4.0 (Chong et al., 2018).

### Meta-analysis of transcriptomics datasets and genome-scale metabolic modeling

In order to understand the regulation of metabolic pathways in PBMCs after seroconversion and T1D progression, genome-scale metabolic models (GEMs) (Bordbar et al., 2012; Sen et al., 2018) of PBMCs were developed. Gene expression or transcriptomics datasets were used to contextualize these models for the P1Ab, P1TD and CTRL groups. Gene expression data of PBMCs was obtained from two, related cohorts, (1) BABYDIET (Ferreira et al., 2014; Hummel and Ziegler, 2011; Schmid et al., 2004), a prospective birth cohort of children in progression to islet autoimmunity and T1D, (2) Diabetes-Genes, Autoimmunity and Prevention (D-GAP) study, a prospective study that recruited children newly-diagnosed with T1D (Ferreira et al., 2014). The longitudinal study settings of these cohorts are similar to the DIPP study design (Knip et al., 2017). The datasets from these studies were downloaded from ArrayExpress (www.ebi.ac.uk; accession number E-MTAB-1724). Expression data from 15 non-progressors (P1Ab), 51 cases of T1D (PT1D), and their controls (CTRL) were selected for genome-scale metabolic modeling (GSMM) (Bordbar et al., 2012; Orth et al., 2010; Sen et al., 2017). In addition, differential expression of genes (DEG) for *P1Ab Vs. CTRL, PT1D Vs. CTRL* and *PT1D Vs. P1Ab* groups was performed. P-values and log2 fold changes were calculated.

A GEM for PBMCs was developed by applying *INIT* algorithm (Agren et al., 2012) on Human Metabolic Reconstruction (HMR 2.0) (Mardinoglu et al., 2014), as a template model. GEMs were contextualized for different conditions using expression datasets. The gene/transcript expression data obtained from PBMCs of PT1D, P1Ab and CTRL was employed to score each reaction of HMR 2.0. Accordingly, the feasibility of a particular reaction(s) to be included or discarded in the draft model was determined. The polar metabolites and the lipidome datasets were used to estimate the confidence score of a metabolite to be included in the draft model. By applying this strategy, condition-specific PBMC models for PT1D, P1Ab and CTRL were developed. Quality control (QC) checks were performed on the draft models (Thiele and Palsson, 2010). In addition, the models were tested for their ability to carry out basic metabolic tasks (Mardinoglu et al., 2013; Mardinoglu et al., 2014). The blocked reactions were removed.

Reporter metabolites (Cakir et al., 2006; Patil and Nielsen, 2005) were predicted by using ‘*reporterMetabolites’* algorithm/function coded in the RAVEN 2.0 suite (Wang et al., 2018). Mixed integer linear programming (MILP) and linear programming (LP) was performed using *‘MOSEK 8’* solver (licensed for the academic user) integrated in the RAVEN 2.0 toolbox. The lower and/or upper bound of an exchange reaction or the uptake rates of a PBMC model were derived from the metabolite concentrations using *‘conc2Rate’* function from COnstraint-Based Reconstruction and Analysis Toolbox (Cobra toolbox v3.0) (Heirendt et al., 2017). Flux Enrichment Analysis (FEA) was performed using *‘FEA’* of Cobra toolbox v3.0. All the simulations were performed in MATLAB 2017b (Mathworks, Inc., Natick, MA, USA).

## DATA ACCESSIBILITY

The metabolomics datasets and the clinical metadata generated in this study were submitted to MetaboLights (Haug et al., 2013), under accession number MTBLS1015. Accompanying clinical metadata was linked to the lipidomics dataset using the ISA-creator package from MetaboLights. The GEMs for PBMCs have been submitted to BioModels (www.ebi.ac.uk/biomodels/), under accession number MODEL1905270001.

## ETHICAL APPROVAL AND INFORMED CONSENT

The ethics and research committee of the participating university and hospital at University of Tampere, Tampere, Finland, approved the study protocol. The study was conducted according to the guidelines in the Declaration of Helsinki. Written informed consent was signed by the parents at the beginning of the study for participation of their children enrolled in the study.

## ACKNOWLEDGEMENTS

We thank the families participating in the DIPP study for making this study possible. We also thank the expert staff of the DIPP study for their excellent work with the participating research families and sample collection. We thank Professor Olli Simell for his important scientific contribution to the DIPP study. We thank Dr. Aidan McGlinchey for assistance with editing the manuscript.

## AUTHOR CONTRIBUTIONS

M.O. and M.K. designed and supervised the study. A.D., T.R., T.L., M.A.L.-B., and E.K., performed the metabolomics experiments, which T.H. supervized. P.S. analyzed the data. S.L assisted with the formulation of statistical design. H.H., J.L, J.T and R.V. contributed to the design of the clinical study. P.S. and M.O. wrote the manuscript. All authors critically reviewed and approved the final manuscript.

