## Supplementary Material for "Metabolic Alterations in Human Peripheral Blood Mononuclear Cells Associate with Progression to Islet Autoimmunity and Type 1 Diabetes"

Partho Sen et al.

### Supplementary Material

**Table S1.** Significantly changed lipids before and after seroconversion (pre vs. post seroconversion analysis).

| Lipids | P1Ab (p-values) | PT1D (p-values) |
|---|---|---|
| Cer(d18:1/24:0) | 0.286 | 0.03 |
| PC(36:3) | 0.036 | 0.309 |
| TG(45:0) | 0.025 | 0.135 |
| TG(47:0) | 0.017 | 0.379 |
| TG(48:0) | 0.041 | 0.013 |
| TG(49:1) | 0.029 | 0.614 |
| TG(50:0) | 0.032 | 0.053 |
| TG(50:1) | 0.016 | 0.007 |
| TG(16:0/16:0/16:0) | 0.302 | 0.011 |
| TG(18:0/18:1/20:4) | 0.7 | 0.03 |
| TG(18:1/12:0/18:1) or TG(18:2/16:0/14:0) | 0.119 | 0.022 |
| TG(18:1/18:1/16:0) | 0.791 | 0.02 |
| TG(18:2/18:1/18:1) | 0.882 | 0.02 |
| TG(48:1) | 0.194 | 0.033 |
| TG(48:3) | 0.9 | 0.042 |
| TG(49:2) | 0.209 | 0.045 |
| TG(50:2) | 0.066 | 0.012 |
| TG(52:2) | 0.305 | 0.018 |
| TG(52:3) | 0.414 | 0.005 |
| TG(53:2) | 0.226 | 0.016 |
| TG(54:6) | 0.932 | 0.007 |

**Table S2.** List of polar metabolites analyzed.

| Metabolite name | Derivitzed metabolite | Retention time | Retention index | Quantification Ion | Qualifier Ion 1 | Qualifier Ion 2 |
|---|---|---|---|---|---|---|
| Alanine | Alanine 3TMS | 17.116 | 1431 | 174.1 | 248.1 | 304.1 |
| Aspartic acid | Aspartic 3TMS | 18.758 | 1517 | 232.1 | 176.1 | 293.1 |
| Citric acid | Citric acid 4TMS | 23.649 | 1825 | 273.1 | 363.1 | 465.1 |
| Cystine | Cystine 4TMS | 30.075 | 2103 | 218.1 | 146.0 | 411.1 |
| Dihydroxyacetoncephosphate | Dihydroxyacetoncephosphate | 20.441 | 1604 | 315.1 | 299.1 | 400.1 |
| Fructose-6-phosphate | Fructose-6-phosphate | 30.075 | 2294 | 315.1 | 217.1 | 459.2 |
| Gluctose-6-phosphate | Gluctose-6-phosphate | 30.189 | 2302 | 387.2 | 357.1 | 471.2 |
| Glutamic acid | Glutamic acid 3TMS | 20.469 | 1605 | 246.1 | 230.1 | 348.1 |
| Lysine | Lysine 4TMS | 25.107 | 1931 | 174.1 | 317.2 | 434.2 |
| Myristic acid | Myristic acid TMS | 24.064 | 1855 | 285.3 | 129.1 | 300.2 |
| Ornithine | Ornithine 3TMS | 22.576 | 1746 | 174.1 | 348.2 | 186.1 |
| Oxoproline | Oxoproline 2TMS | 18.747 | 1516 | 156.1 | 230.1 | 258.1 |
| Palmitic acid | Palmitic acid TMS | 26.859 | 2059 | 313.2 | 132.0 | 328.3 |
| Pentanedioic acid | Pentanedioic acid(imino) 2TMS | 19.724 | 1567 | 147.1 | 198.0 | 304.1 |
| Phenylalanine | Phenylalanine 3TMS | 20.53 | 1608 | 218.1 | 192.1 | 266.1 |
| Phosphoenolpyruvate | Phosphoenolpyruvate | 20.121 | 1587 | 369.1 | 299.1 | 384.0 |
| Proline | Proline 2TMS | 14.637 | 1303 | 142.1 | 216.1 | 244.1 |
| Serine | Serine 3TMS | 15.845 | 1366 | 204.1 | 218.1 | 278.1 |
| Stearic acid | Stearic acid TMS | 30.189 | 2302 | 341.3 | 117.0 | 356.3 |
| Threonine | Threonine 3TMS | 16.339 | 1391 | 218.1 | 117.1 | 291.1 |
| Tyrosine | Tyrosine 3TMS | 25.331 | 1947 | 218.1 | 280.1 | 382.2 |
| Unknown BCAA 1 | BCAA 1 | 11.897 | 1161 | 86.1 | 75.0 | 188.0 |
| Unknown BCAA 2 | BCAA 2 | 12.314 | 1183 | 86.1 | 75.0 | 188.0 |
| Unknown BCAA 3 | BCAA 3 | 12.734 | 1204 | 86.1 | 75.0 | 188.0 |
| Valine | Valine 2TMS | 10.651 | 1096 | 144.1 | 100.1 | 218.1 |

**Figure S1.** Factors and sources of variation in the metabolomics datasets, displaying a density plot of the metabolite-wise marginal $R^2$ values for the factors. Effects of different factors such as age, sex and study group status on lipidomics data were evaluated. The data were centered to zero mean and unit variance. The relative contribution of each factor (experimental variable) to the total variance in the dataset was estimated by fitting the linear model regression model, where the normalized intensities of metabolites regressed to the factor of interest, and thereby estimating the median marginal coefficients ($R^2$). This analysis was performed using package 'scater'.

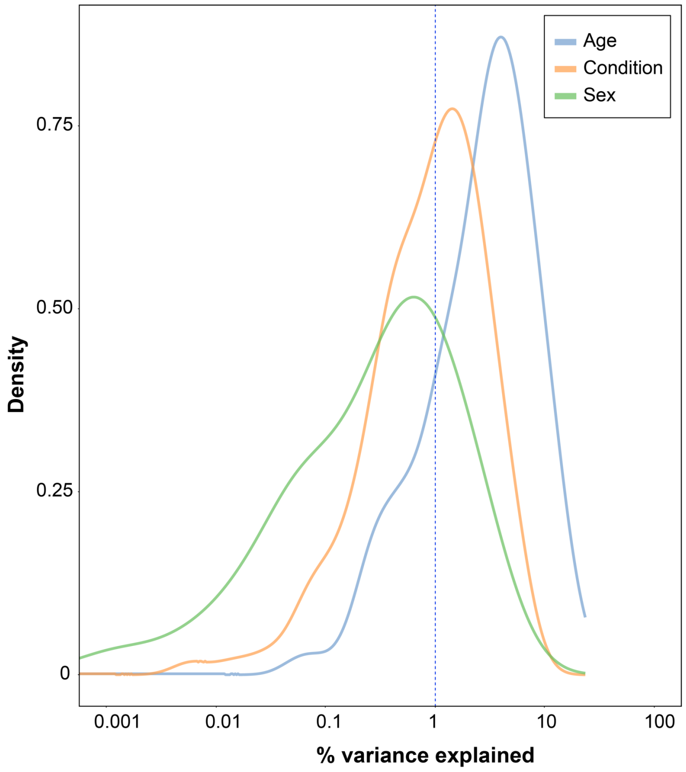

**Figure S2.** Selected lipid profiles that are significantly altered in CTRL, P1Ab and PT1D groups, during the 36-months follow-up. A-F) Time-course profiles of sphingomyelin (SMs), cholesterol esters (CEs), ceramides (Cer), lysophosphatidylcholine (LPCs), phosphatidylcholine (PCs), and triacylglycerides (TGs) are shown.

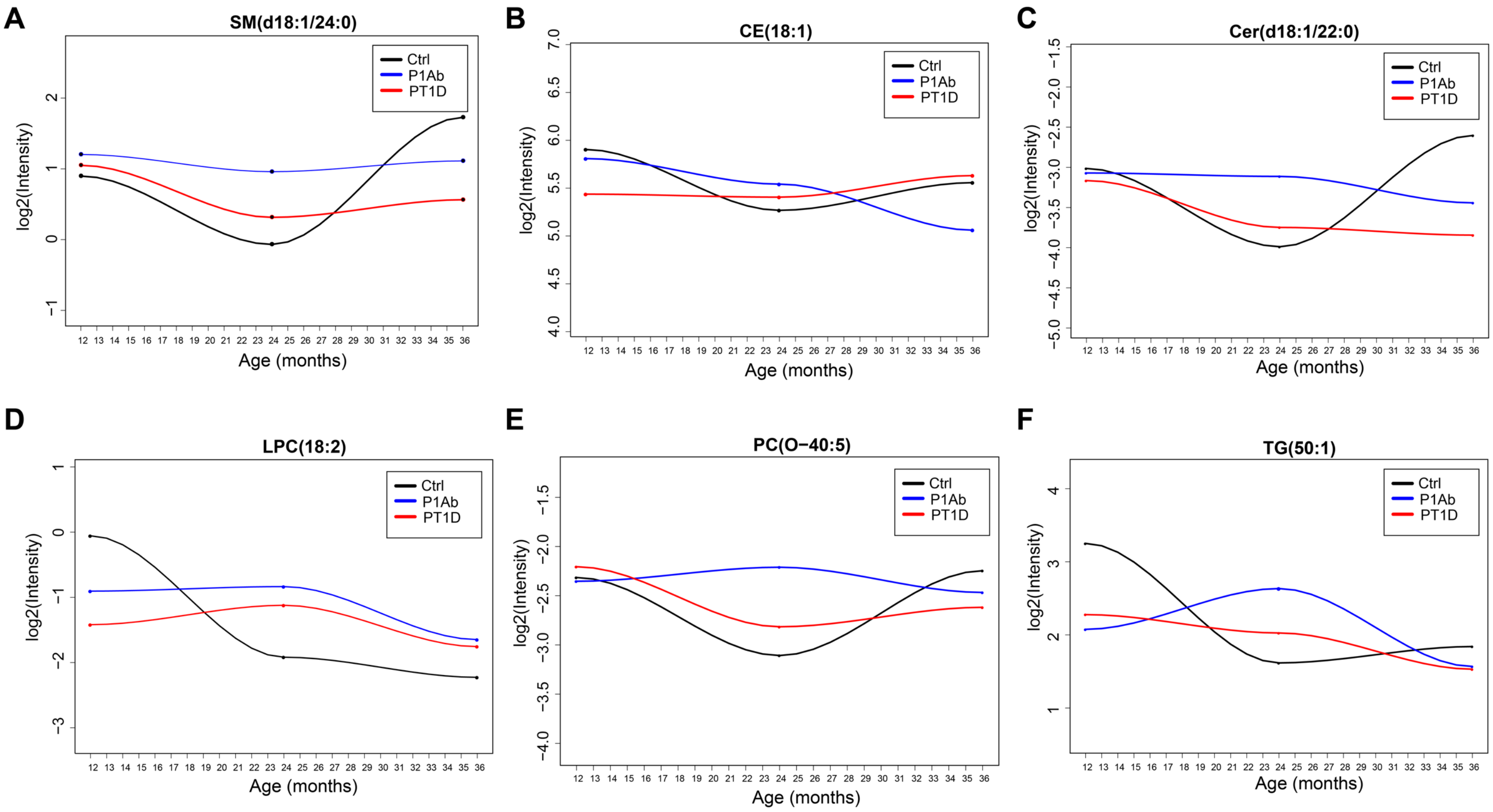

**Figure S3.** log intensities of the total lipids before (light blue) and after (light gray) the seroconversion (SC), in P1Ab and PT1D groups. Significant down-regulation in the intensities of the total lipids in the PBMCs were observed after seroconversion (SC), in P1Ab (median age of SC, 24 months) (p=5.581e-05) and PT1D (median age of SC, 14 months) (p=9.803-06).

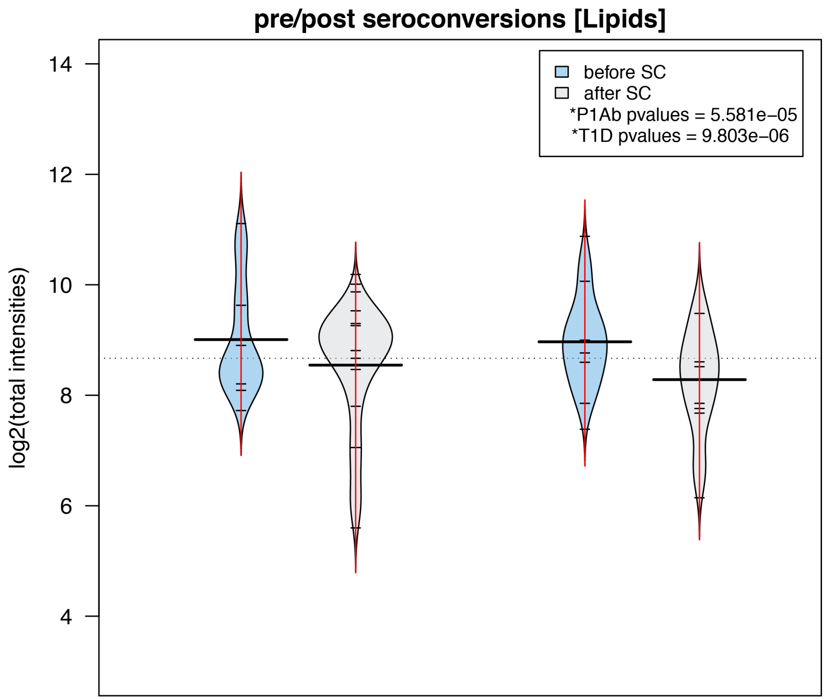

**Figure S4.** Selected profiles of polar metabolites that are significantly altered in CTRL, P1Ab, and PT1D groups, during the 36-months follow-up. A-F) Time-course profiles of palmitic acid, aspartic acid, citric acid, myristic acid, phenylalanine, and serine are shown.

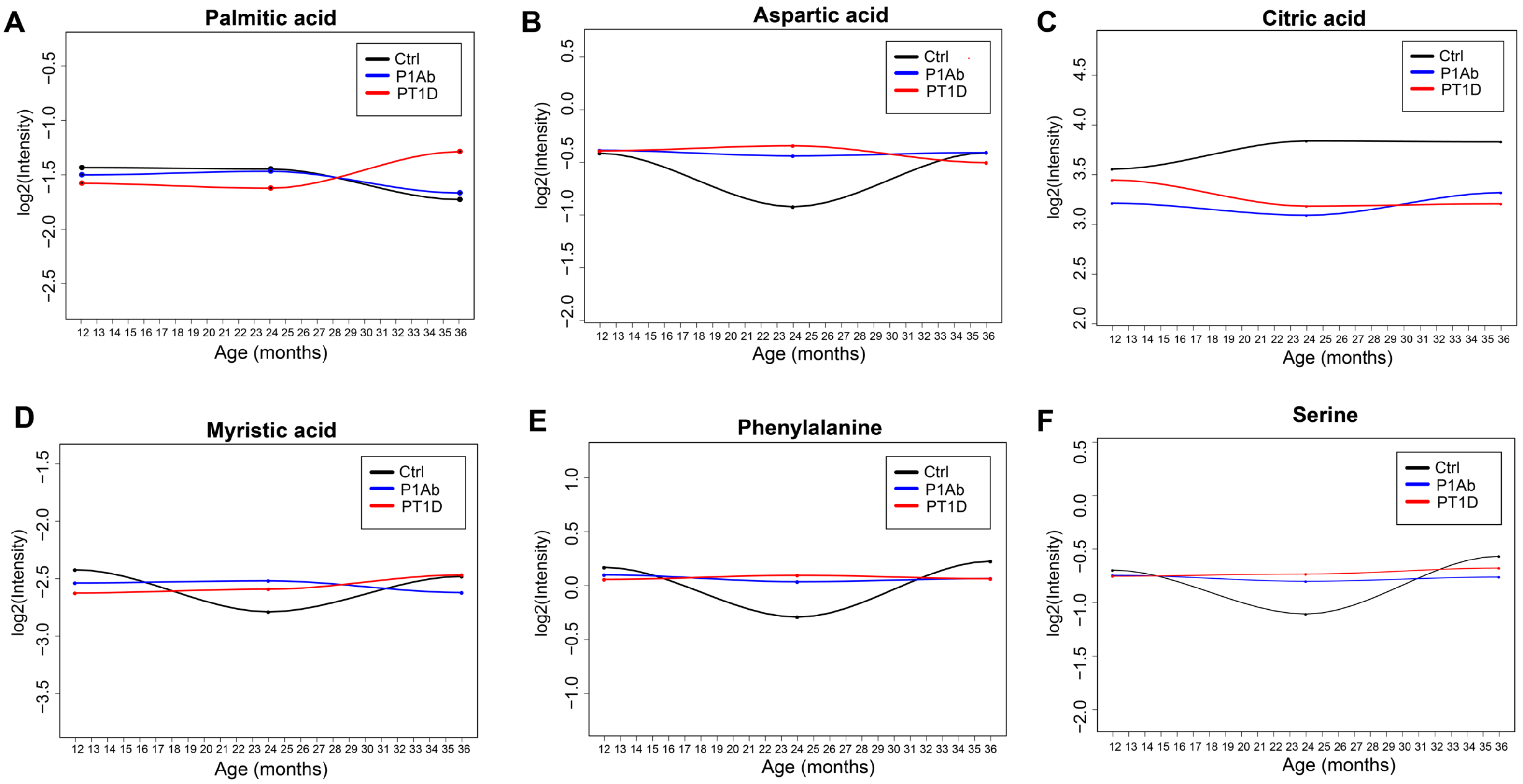

**Figure S5.** Heatmap showing Spearman's correlation between plasma and cellular (PBMCs) metabolite intensities in P1Ab at 12 and 36-months of follow-up. Red, blue and white color suggests positive, inverse and no correlation respectively.

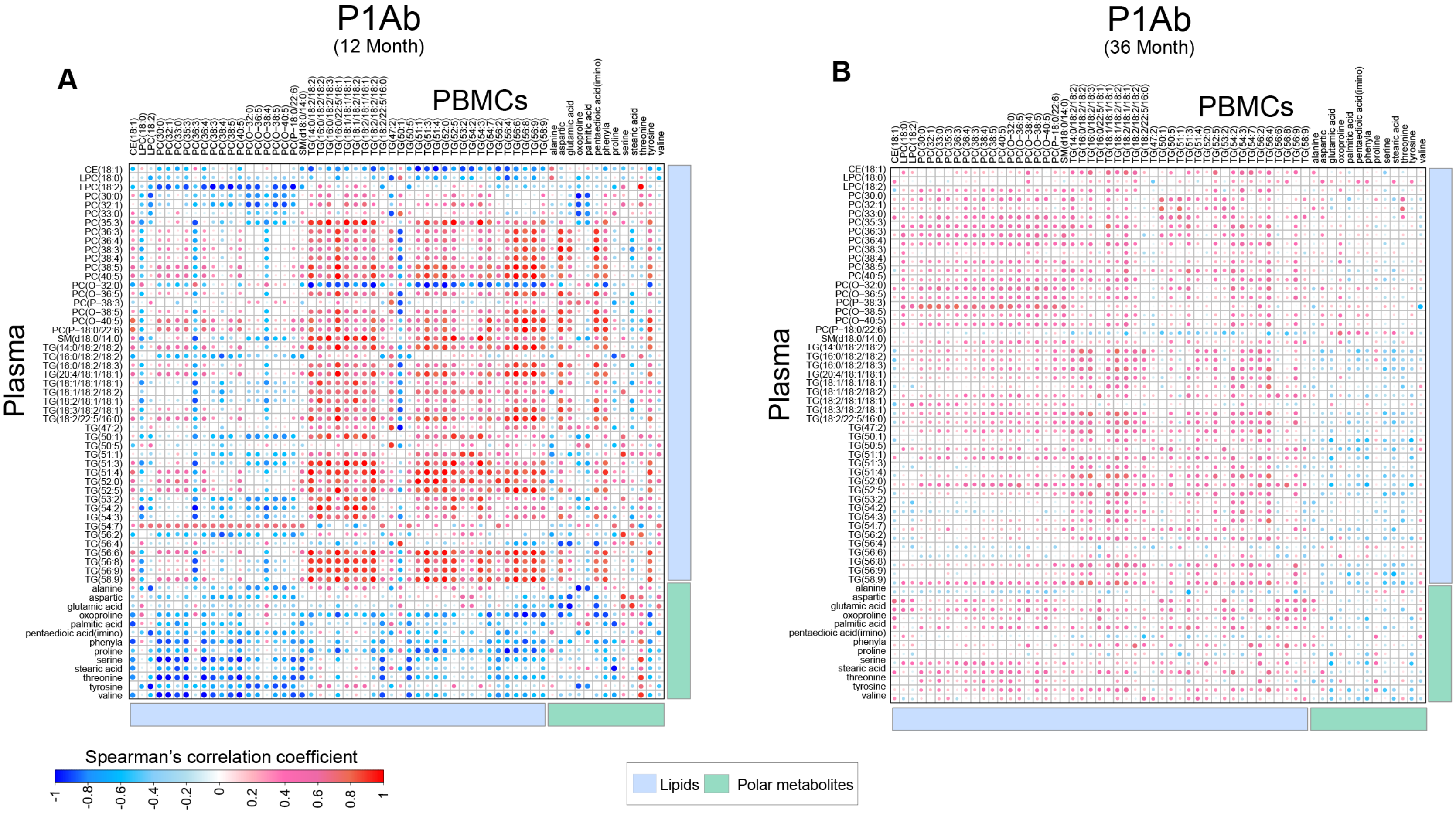

**Figure S6.** Heatmap showing Spearman's correlation between plasma and cellular (PBMCs) metabolite intensities in CTRL and PT1D at 36-months of follow-up. Red, blue and white color suggests positive, inverse and no correlation respectively.

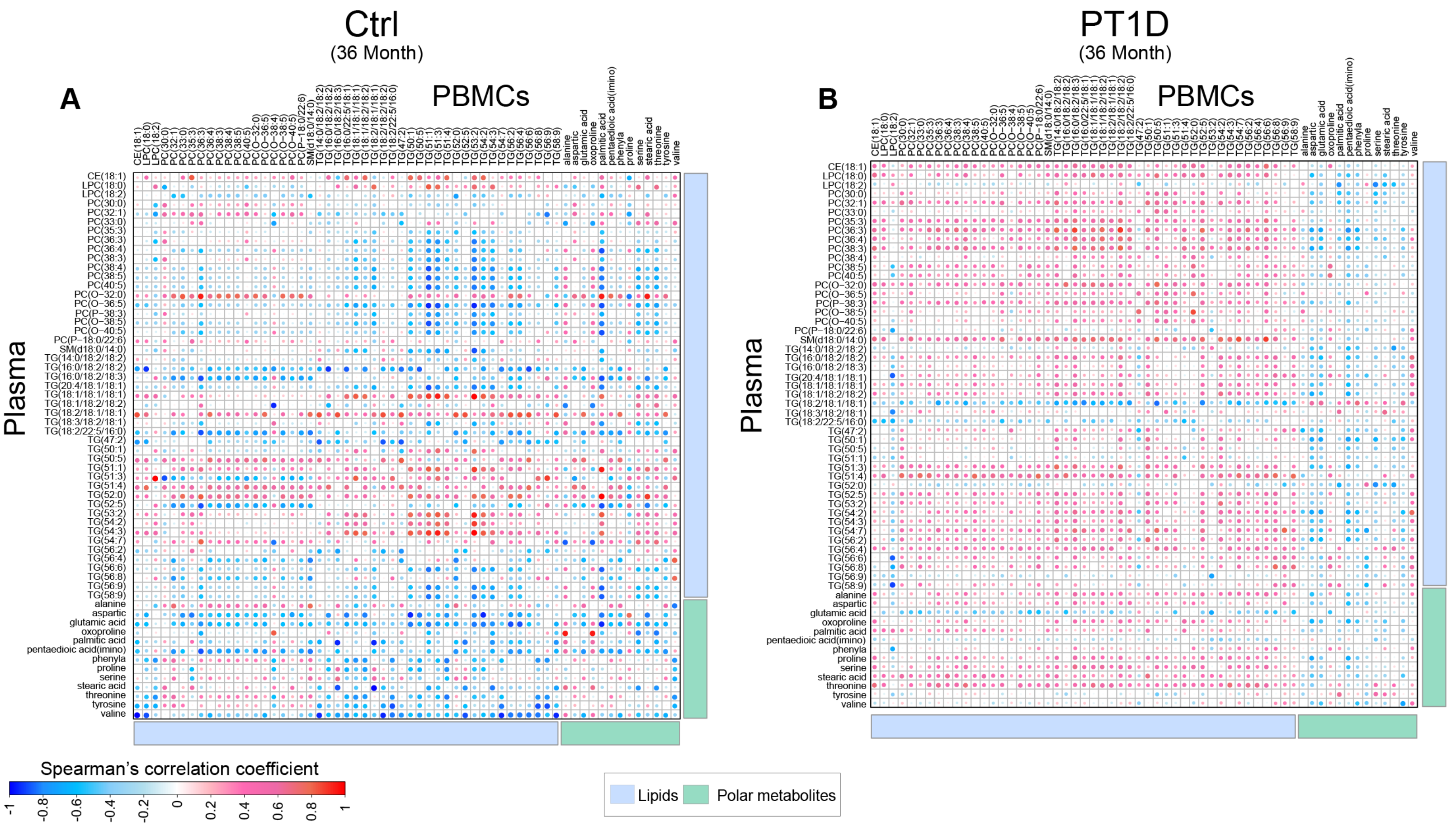

**Figure S7.** Genome-scale metabolic models for PBMCs. The total numbers of genes, metabolites and reactions included in the PBMC models for CTRL, P1Ab and PT1D, developed in the study is shown.

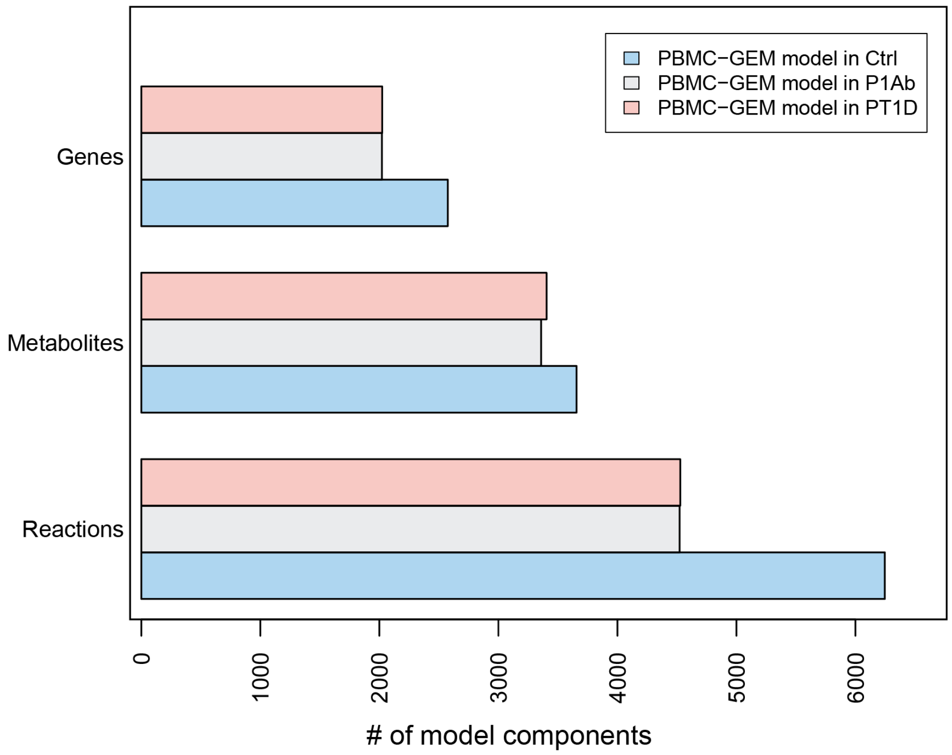


**Figure.S10**

**Figure S8.** Reporter metabolites of PBMCs, based on genome-scale metabolic modeling, that were significantly changed (FDR < 0.05) in PT1D vs. CTRL.

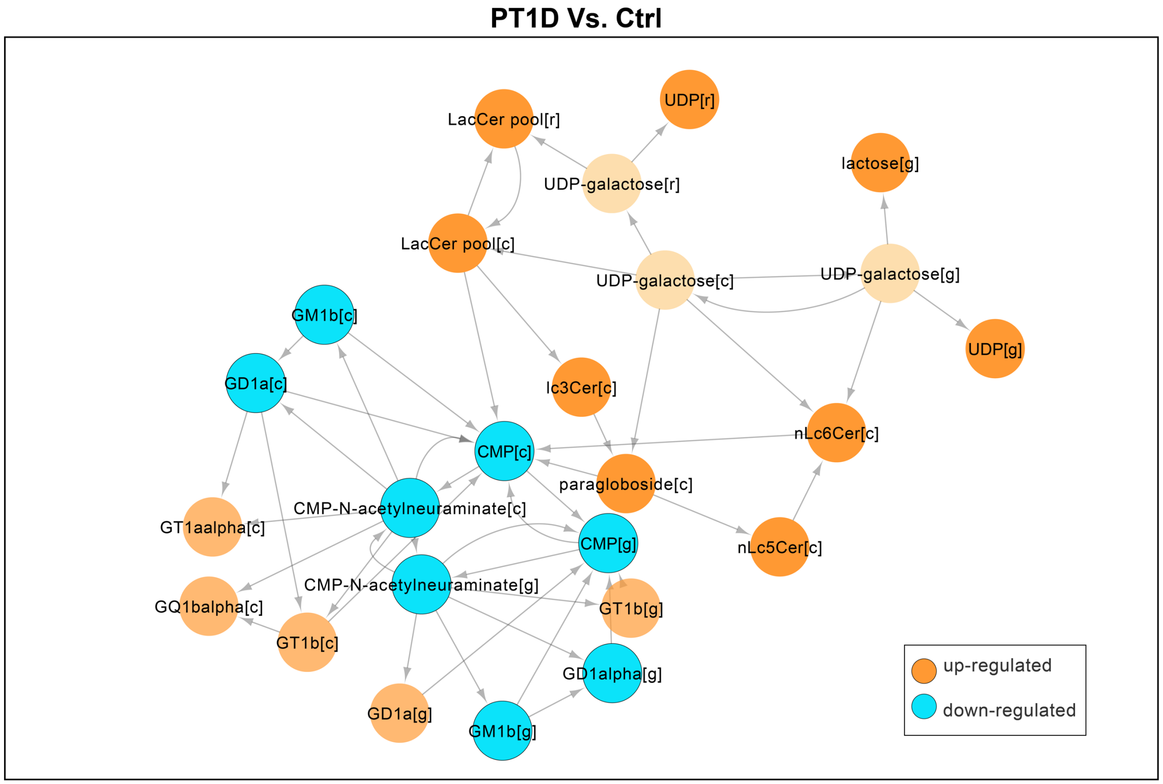

**Figure S9.** The plot shows reporter metabolites of PBMCs that were significantly changed (FDR < 0.05) in P1Ab vs. CTRL.

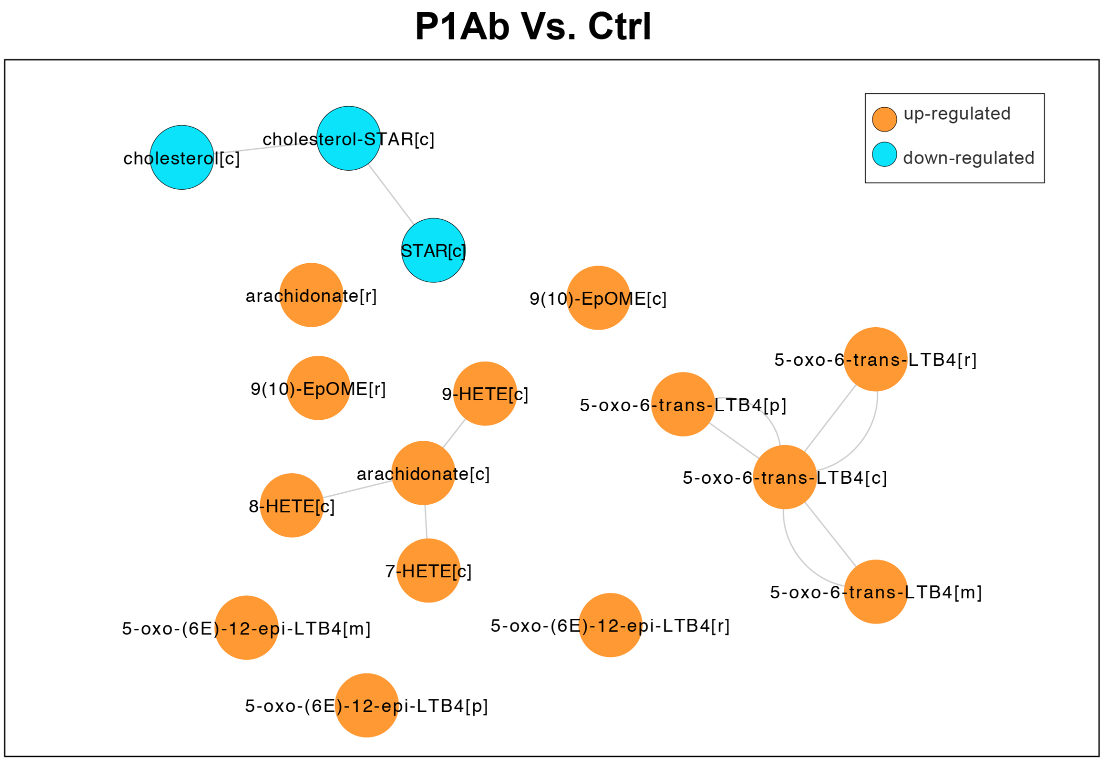

**Figure S10.** Flux enrichments in the metabolic subsystems. Stacked bar plots showing cumulative (-$\log_{10}$) q-values of the enriched subsystems. Enrichments of metabolic subsystems in PBMCs models for T1D progressors and nonprogressors when, A) production of glucosylceramide is maximized, B) production of digalactosylceramide from D-galactosyl-N-acylsphingosine is maximized.

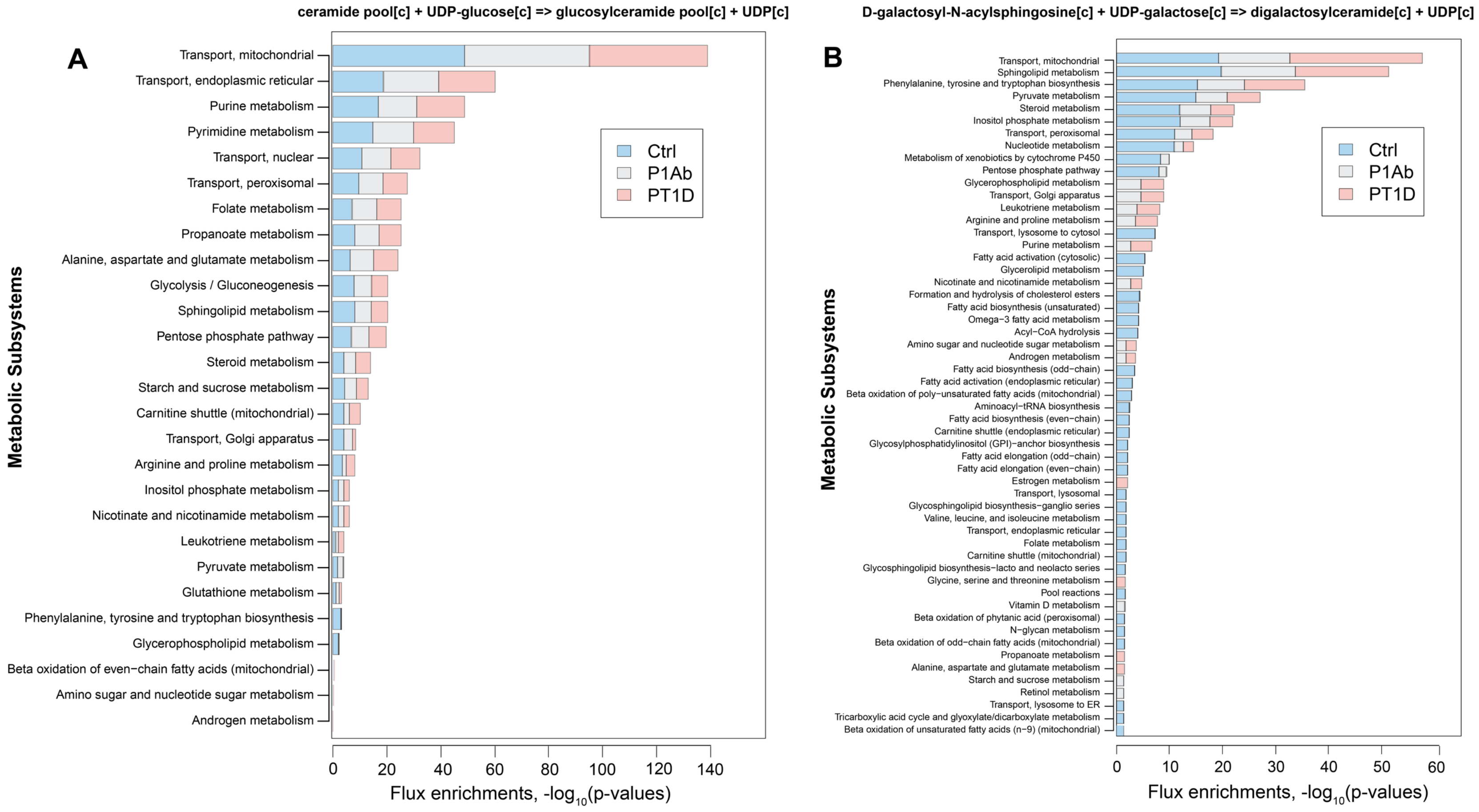

**Figure S11.** Bar plot showing the number of subjects included in the, (i) **CTRL group**: children who remained autoantibody negative (*Ab*⁻) during the follow-up. (ii) **P1Ab group**: children who developed at least one islet autoantibody but were not diagnosed with Type 1 diabetes (T1D), during the follow-up. (iii) **PT1D group**: comprises of children seroconverted to multiple islet autoantibodies and later developed T1D. The subjects are ordered by their age groups.

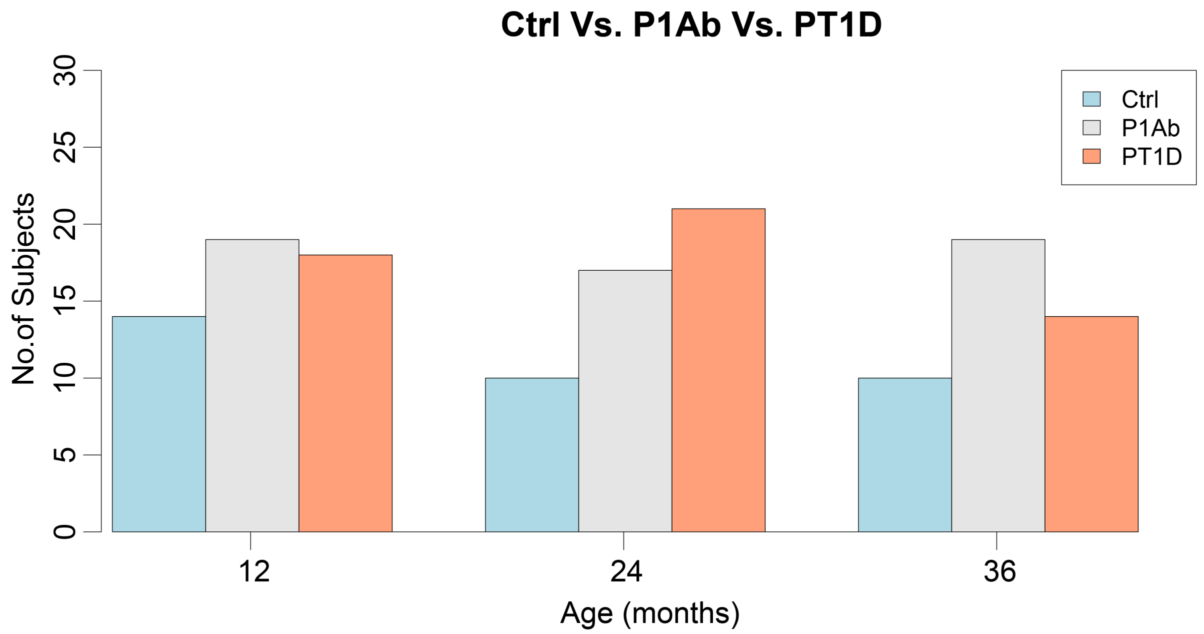

**Figure S12.** Log normalized intensities of the lipids measured in CTRL, P1Ab and PT1D groups. The subjects are colored and ordered by their age groups. The outliers are marked by grey dots.

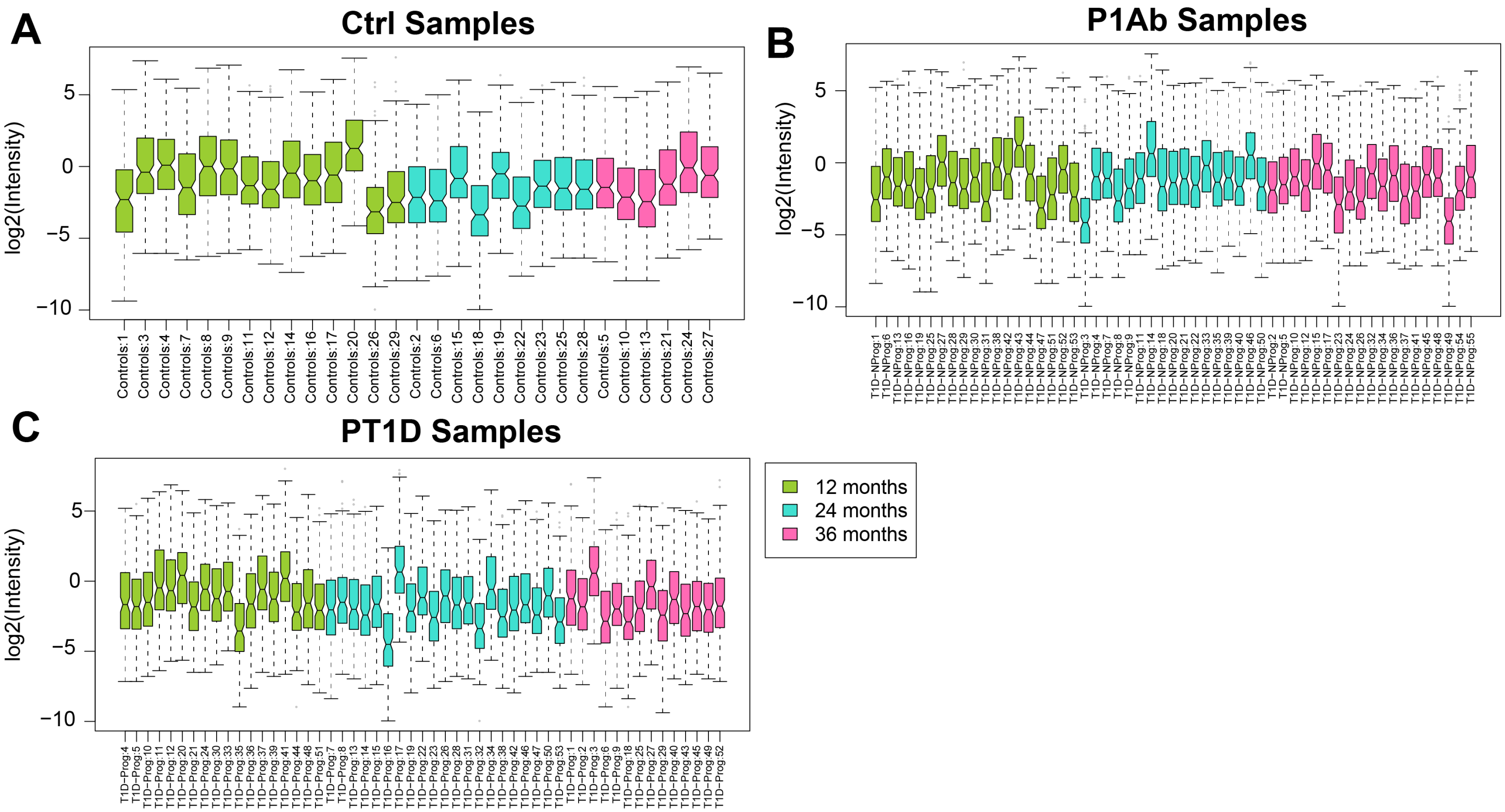

**Figure S13.** Log normalized intensities of the polar metabolites measured in CTRL, P1Ab and PT1D groups. The subjects are colored and ordered by their age groups. The outliers are marked by grey dots.

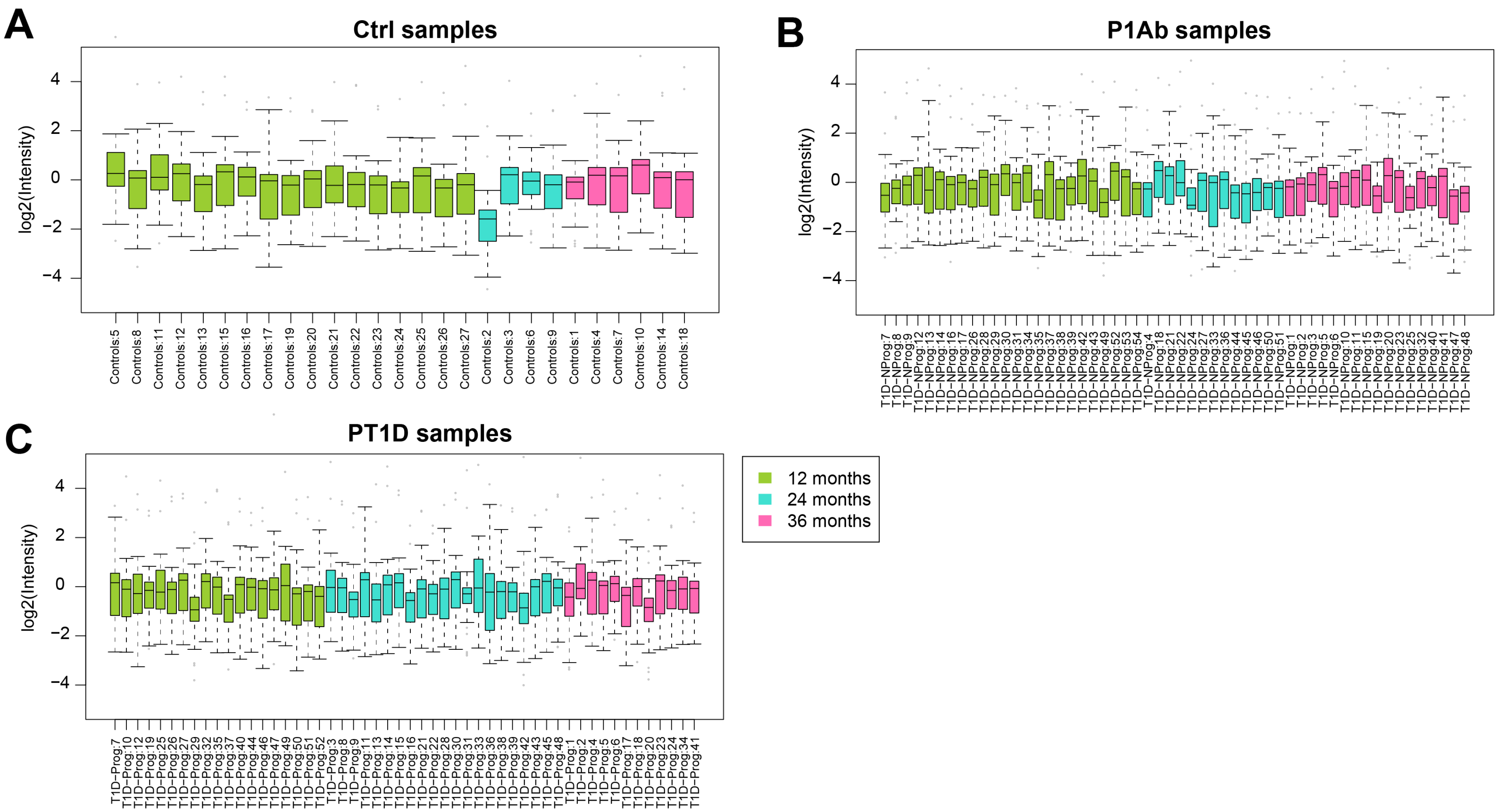
